# A local composition of peptidoglycan drives the division site selection by MapZ in *Streptococcus pneumoniae*

**DOI:** 10.1101/2025.04.08.647842

**Authors:** Adrien Ducret, Céline Freton, Christophe Grangeasse

**Affiliations:** Molecular Microbiology and Structural Biochemistry, UMR 5086, Université de Lyon, CNRS, Lyon, France

**Keywords:** Streptococcus pneumoniae, MapZ, division site selection, peptidoglycan, bacterial cell division

## Abstract

Accurate division site placement is essential for bacterial cells to produce viable daughter cells with proper size and appropriate functional features. In the opportunistic pathogen bacterium *Streptococcus pneumoniae*, the positioning of the division site has been shown to depend on both the protein MapZ and chromosome segregation. However, the nature of this interplay and the molecular determinants guiding division site localization remained unclear. Here we demonstrated that the division site is positioned at the cell equator, the widest part of the cell body, rather than at mid-cell. In addition, we observed that the localization of MapZ and/or the divisome remain unaffected even in the absence of properly segregated DNA, indicating that chromosome segregation does not contribute to division site selection. Our findings further reveal that MapZ localization depends on the activities of two PG hydrolases DacA and DacB, whose sequential recruitment to the division site during early PG synthesis drives the formation of a distinctive PG signature required for MapZ binding. These results support a model in which MapZ identifies the division site by recognizing a specific PG composition produced only during the early stages of cell division. This PG composition becomes enriched at the cell equators which will eventually serve as the division site of the daughter cells.

## Introduction

Bacterial cell division involves the formation of a new cell wall to divide a single bacterium into two daughter cells. Cell division in bacteria is primarily driven by the protein FtsZ, which forms a ring-like structure known as the Z-ring at the site of division^1^. This Z-ring serves as a scaffold for the recruitment of additional proteins necessary for cell constriction and the synthesis of the cross-wall, thereby forming a division machinery known as the divisome. To ensure the accurate partitioning of cellular components and the generation of viable offspring, the placement of the site of division must be tightly regulated. The placement of the divisome, and more specifically the formation of the Z ring, is influenced by multiple factors, including the localization of specific proteins, the positioning of the bacterial chromosome, and signaling mechanisms within the cell^2^. In well-studied rod-shaped model organisms, such as *Bacillus subtilis* and *Escherichia coli*, the placement of the division septum at mid-cell is controlled by the well-known nucleoid occlusion and the Min systems^3,4^. Both systems are negative regulators, which inhibit the assembly of the Z ring at inappropriate locations, such as over the nucleoid or near cell poles, and promote its localization at mid-cell. In the past decade, studies on different bacterial models that exhibit distinct developmental programs, cell shape or undergo asymmetric division led to the discovery of distinctive and unique systems that exert either a positive or a negative influence on the positioning of FtsZ at the division site. For instance, the placement of the Z-ring at the cell center of the α-proteobacterium *Caulobacter crescentus* is coordinated with chromosome segregation by the protein MipZ that directly interferes with the polymerization of FtsZ^5^. On the other hand, the protein SsgB in *Streptomyces coelicolor* recruits and enhances the polymerization of FtsZ during sporulation^6^, while the ParA-like protein PomZ localizes prior to FtsZ at the division site and reinforces the stability of the Z-ring in *Myxococcus xanthus*^7^. More recently, the protein PcdA in *Staphylococcus aureus* interacts with the structural protein, DivIVA to recruit unpolymerized FtsZ and promote the assembly of the Z-ring along the proper cell division plane^8^. However, the exact mechanisms driving the localization of these proteins and how they coordinate and establish the division site in possible conjunction with the nucleoid is still an active area of research and likely involve multiple regulatory pathways. This is especially true for the placement of the division site in the opportunistic pathogen *Streptococcus pneumoniae*, which is proposed to rely both on the protein MapZ^9,10^ and chromosome segregation^11^.

MapZ is a single-spanning membrane protein with a cytoplasmic domain that interacts directly with FtsZ and an extracellular domain that interacts with the peptidoglycan (PG), the major constituent of the cell envelope^12^. MapZ forms a ring-like structure at mid-cell prior FtsZ and then recruits FtsZ, FtsA and EzrA to guide septum placement^9,10,13,14^. Following the formation of the Z-ring, duplicate MapZ rings migrate away from the septum, beaconing the future division sites as the cell elongates. Upon division, the two MapZ ring-like structures could serve as nanotracks guiding and constraining the treadmilling of FtsZ filaments moving from the division septum to the new sites of division of daughter cells^13,15^. Although nonessential, the loss of MapZ results in asymmetric cell division from mis-localized Z rings: FtsZ-rings no longer positioned properly and the angle relative to the cell long axis is incorrect^9,10^. Alternatively, both the origin regions (*OriC*) of the replicating chromosome also localize near the future mid-cell of the daughter cells prior FtsZ and MapZ and were consequently proposed to also contribute to the placement of the division site in *S. pneumoniae*^11^. Moreover, when the longitudinal chromosomal organization is affected by poisoning DNA decatenation or by mutating the condensin SMC, the re-localization of MapZ and FtsZ at the future site of division is also affected, causing irregular cell division^11^. While MapZ and the nucleoid may perform distinct roles in the selection of the division site, this implies a close relationship between the role of MapZ and chromosome segregation and the selection of division sites in the pneumococcus. However, their interplay, the molecular determinants and the exact mechanisms driving the localization of the division site remain unknown. Here, we demonstrated that the formation of the division septum does not occur at the future mid-cell of the daughter cells but rather at their cell equators, the widest part of their cell body. By looking at the sub cellular localization of MapZ and *OriC*, we observed that only MapZ localizes at the cell equators throughout the cell cycle while *OriC* was more dispersed and longitudinally offset. We further demonstrated that the localization of MapZ and the divisome, at the cell equators were not affected even in the absence of properly segregated DNA showing that *OriC* or the nucleoid are not involved in the selection of the division site in *S. pneumoniae* as suggested before^11^. We then showed that the localization of MapZ at the cell equators was compromised in the absence of the DacA and/or DacB, two carboxypeptidases that consecutively degrade the stem-peptide involved in the reticulation of the PG. Furthermore, our findings indicate that MapZ specifically recognizes a distinctive signature, involving tetrapeptides which could be produced by the sequential recruitment of DacA and DacB at the division site. Together, these findings reveal that the selection of the division site by MapZ in *S. pneumoniae* is driven by a local PG composition at the cell equators.

## Results

### The site of division does not localize at mid-cell in *S. pneumoniae*

It is widely accepted that cell division in *S. pneumoniae* occurs by the formation of a division septum at mid-cell^16^. To assess this, we examined the longitudinal localization of FtsA-mKate as a proxy for the divisome in nascent cells. Strikingly, we found that their position was offset from mid-cell (Fig. 1A, p<0.0001, unpaired t-test), with at least 9±2% of Z-rings located more than 0.15 µm away. This suggested that division septum formation in *S. pneumoniae* did not strictly occur at the mid-cell. Interestingly, those rings were consistently observed at the widest part of asymmetrical cells with two poles differing significantly in size and shape (Fig. 1B&C). Unlike rod-shaped bacteria such as *E. coli* and *B. subtilis*, which incorporate new PG along their lateral walls, *S. pneumoniae* incorporates new material exclusively at the division site, forming a new cell pole at each division cycle^17^. Consequently, the daughter cells always consist of a newly synthetized pole and an old pole inherited from the mother cell. The formation of asymmetrical cell is thus likely driven by inherent regulatory variations between constriction and elongation across successive division cycles (Extended Data Fig. 1). Importantly, this also infers that the widest part of the cell, also known as the cell equators, always lies at the junction between the newly synthetized pole and the old pole inherited from the mother cell. To determine if the formation of the division septum occurs specifically at this interface during the cell cycle, we examined the localization of FtsA-mKate along with that of the cell equators using PG labeling at the single-cell level using time-lapse microscopy (Fig. 1D-F and Video 1). In this experiment, PG of the whole cell was labeled by long pulse of the fluorescent D-amino acid (FDAA) followed by a chased experiment performed under the microscope. In the early steps of the cell cycle, FtsA-mKate formed a ring at the equator while the whole cell periphery is labelled with FDAA. As the cell elongates, the formation of non-stained areas reflected the insertion of new material at the active division site forming the new cell-halves (unstained) while the old cell-halves (stained) are pushed apart (Fig. 1D-F and Video 1). Consequently, the interface between the new and old cell wall material keeps moving away from the active division site as the cell elongates, marking the cell equators of the future daughter cells throughout the cell cycle. While most of the FtsA-mKate signal remains at the active division site during cell cycle progression, a fraction of FtsA-mKate re-localized at the equators of the future daughter cells (Fig. 1D-F). This location will eventually become the future sites of division once the division is completed. Altogether these results show that the future sites of division are not located at the future mid-cell of the daughter cells but rather at their cell equators.

**Figure 1:**
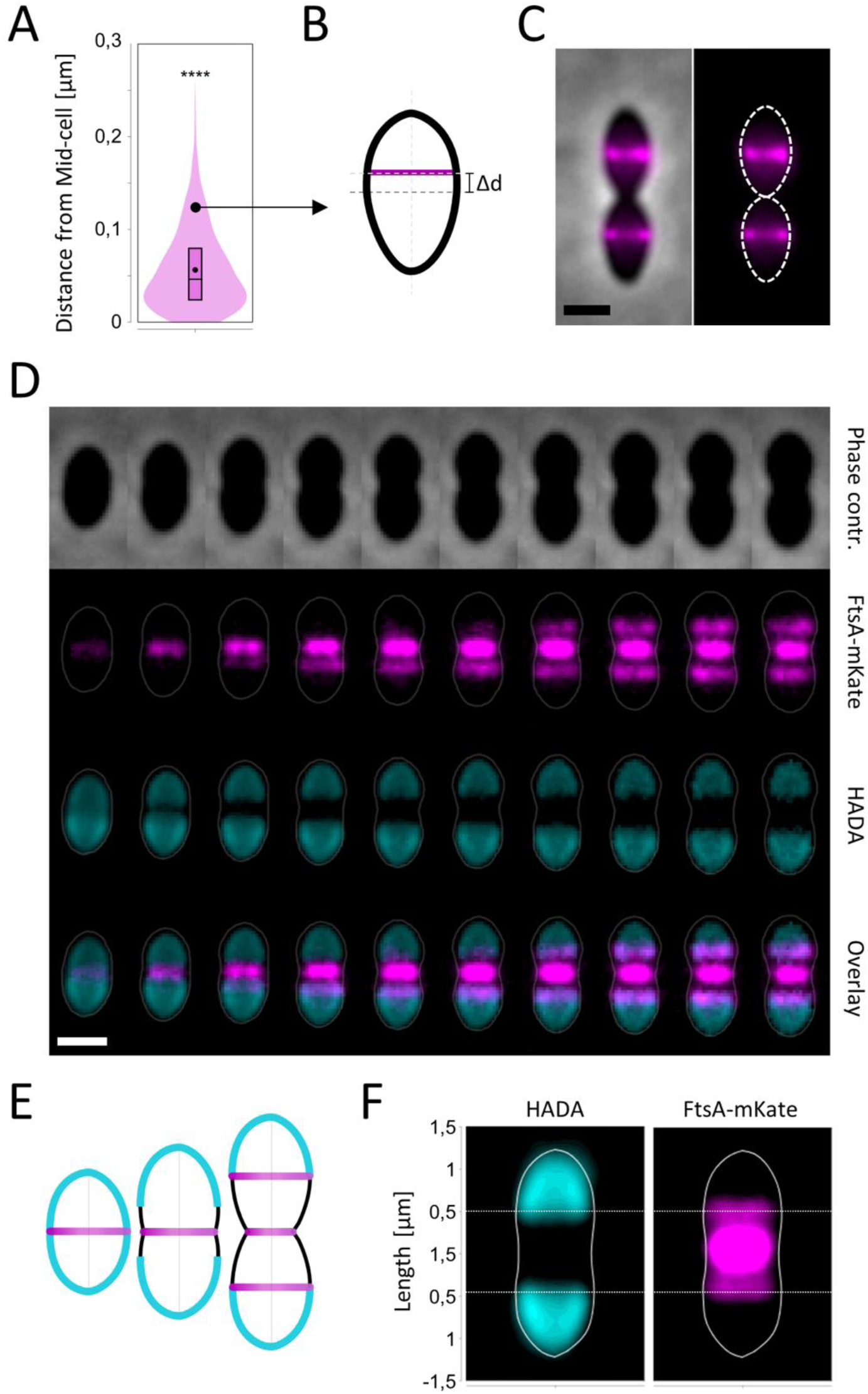
The site of division does not localize at mid-cell in *S. pneumoniae.* Violin plots representing the distribution of the distance (Δd) between the longitudinal position of the FtsA-ring and mid-cell in a population of cells harvested at the mid-exponential phase of growth in C+Y media at 37°C (n=2510, r=3, p<0.0001 – Unpaired T-test). (B) Schematic of the contour of a cell where the longitudinal position of the FtsA-ring is significantly offset relative to mid-cell (Δd) as shown in (C). The cell medial axis is represented in light gray, the transversal axis at mid-cell in dark gray and the transversal FtsA-ring in magenta. (C) A representative image showing the localization of mKate2-FtsA in two asymmetrical daughter cells. The mKate-channel, phase contrast and merged images are shown. The dash line on the merged image represents the cell contour retrieved from MicrobeJ. The scale bar represents 1 µm. (D) A representative montage of images showing the localization of mKate2-FtsA and the interface between the new and the old poles (aka cell equators) as the cell elongates at the single cell level by time-lapse microscopy. The cell equators can be visualized using a long pulse of the fluorescent D-amino acid (FDAA) followed by a chased experiment performed under the microscope. Pictures were taken every 3 min. The phase contrast, mKate-channel, the HADA-channel and the merged between the mKate and the HADA channels images are shown. The scale bar represents 1 µm. (E) Schematic representation of the contour of a cell labeled by long pulse of HADA followed by a chased experiment performed under the microscope and the localization of the divisome (magenta). The newly synthetized pole (black) and the old poles inherited from the mother cell (light blue) are shown. (F) Averaged sub-cellular localization of mKate2-FtsA and the interface between the new and the old poles (n=37).

### MapZ, but not *OriC*, remains localized at the cell equators throughout the cell cycle

The chromosomal origin of replication (*OriC*), is proposed to localize at the future mid-cell before MapZ, implying a potential role in positioning the division site in *S. pneumoniae*^11^. Since we demonstrated that the division site is not necessarily at mid-cell, we reassessed the localization of *OriC* and wondered whether it could contribute to the localization of the future site of division. To test this, we first looked at the sub cellular localization of the divisome, the equators and either MapZ or *OriC* at the population level. To this end, we constructed two strains expressing FtsA-mKate as a proxy for the divisome, and either mGfp-MapZ or ParB-mGfp which binds to *OriC* as described in^11^ and thereby allows to visualize the origin of replication. In addition, we developed a multipeak-fitting approach in MicrobeJ^18^ to improve the detection of near FtsA and MapZ rings, or near ParB-mGfp foci during the initial steps of the division process^19^ (Extended Data Fig. 2A). Then, the position of the rings or the foci on the medial axis of the cells was plotted against the length of the corresponding cells on a density histogram (Fig. 2A). In the early steps of the cell cycle, FtsA-mCherry and mGfp-MapZ positioned at the equators of nascent cells. While most of the FtsA-mCherry signal remains at the active division site, mGfp-MapZ rings split into two rings that migrated with the cell equators as the cell elongate (Fig. 2A). During cell cycle progression, FtsA-mCherry re-localized with mGfp-MapZ at the equators of the future daughter cells. By contrast, the duplication of the ParB-mGfp focus was followed by rapid segregation of the two foci into the old cell-half of cells where they remain as the cell elongate (Fig. 2B). In addition, we observed that the distribution of the localization of *OriC* was more dispersed than that observed for FtsA-mCherry and mGfp-MapZ (Fig. 2C&D). Indeed, the ParB-mGfp foci were in average offset (toward the old pole) relative to the equators (Fig. 2D). To further support this observation, we examined the cellular localization of the divisome, the equators, and either MapZ or OriC at the single-cell level (Extended Data Fig. 2B). While FtsA-mKate and mGfp-MapZ strictly co-localized at the cell equators, ParB-mGfp foci kept oscillating on both sides of the cell equators and were longitudinally offset relative to the future cell division septum (Extended Data Fig. 2B). Consequently, *OriC* does not position systematically at the division site nor does it position there prior to MapZ.

**Figure 2:**
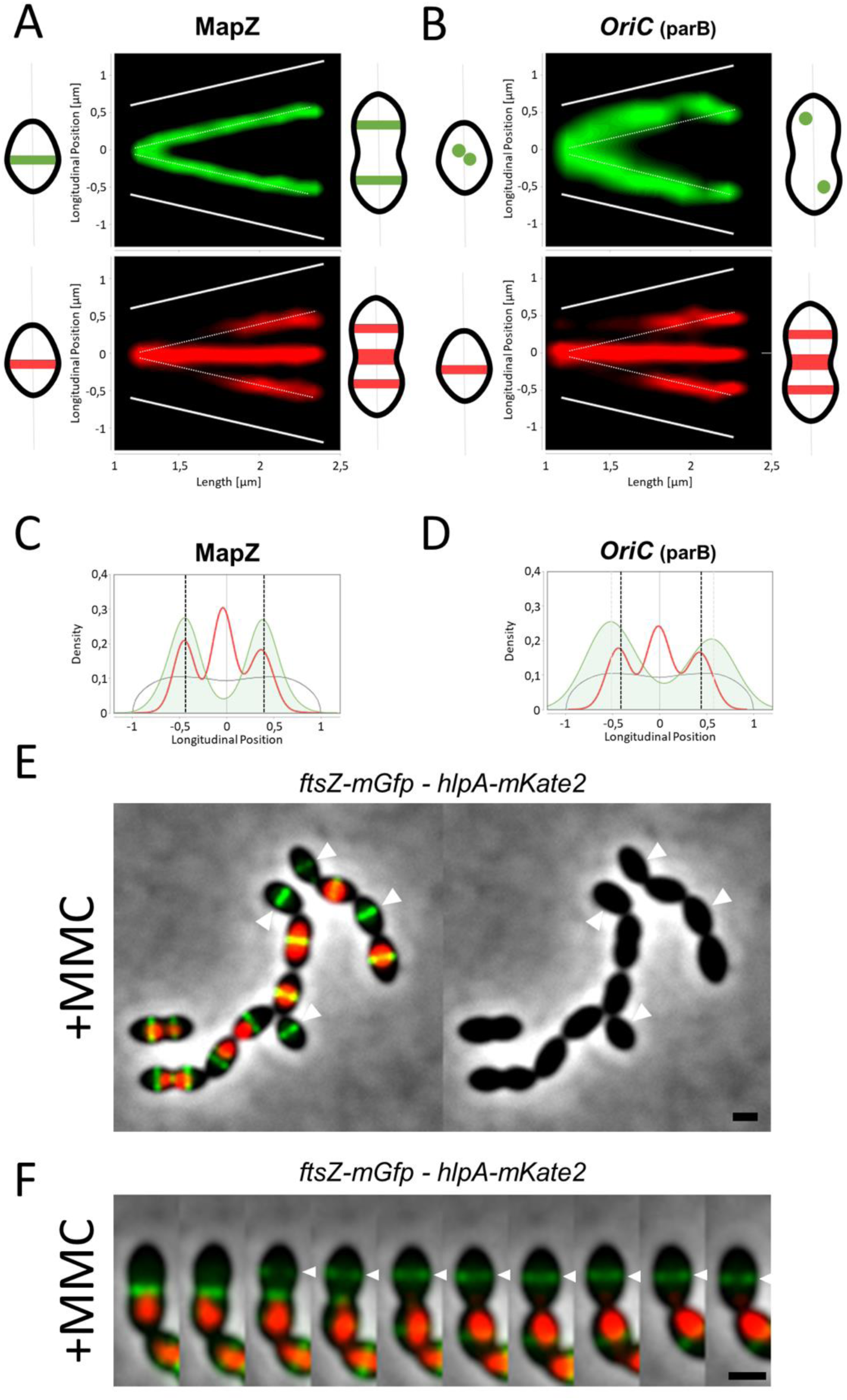
MapZ, but not Ori, localizes at the cell equators throughout the cell cycle. (A-B) The longitudinal position of FtsA-mKate and mGfp-MapZ rings (A) or ParB-mGfp foci detected by MicrobeJ as a function of the cell length, shown as heatmaps. The color saturation represents the density of the localizations. The white dashed lines represent the cell equators and the solid lines the tip of the poles. The data represent a total of 2517 mGfp-localizations/2989 mKate-localizations. A schematic of a representative localization of FtsA-mKate, mGfp-MapZ ring(s) and ParB-mGfp foci in a new born and pre-divisional cell are shown on both sides of each graph for better clarity. (C-D) The relative longitudinal position of FtsA-mKate (red line) and mGfp-MapZ rings (green line;C) or ParB-mGfp foci (green line; D) shown as density plots for cells with a cell length ranging from 1.8 µm to 2 µm. The averaged shapes for each condition are shown in gray. (E) Representative image showing the localization of FtsZ-mGfp at the cell equators in cells whose chromosome (detected using a HlpA-mKate2 fusion) was either not or only partially segregated after treatment with a sub-lethal dose of MMC (7.5ng/ml). The white triangles highlight the FtsZ rings at the cell equators of cells where the nucleoid is not detectable. The merged of the phase contrast, the mKate-channel and the gfp-channel images, and the phase contrast images are shown. The scale bar represents 1 µm. (F) A representative montage of images showing the localization of FtsZ-mGfp and HlpA-mKate (as a proxy for the nucleoid) as the cell elongates at the single cell level by time-lapse microscopy. The white triangle highlights the re-localization of the Z-ring from the active division site to the future division site of division of a daughter cell without a detectable nucleoid. Pictures were taken every 2 min. The merged between the mKate, the mGfp channels and the phase contrast images are shown. The scale bar represents 1 µm.

To check that *OriC* or the nucleoid may play a role into the placement of the division site, we looked at the localization of either FtsZ-mGfp, FtsA-mGfp or mGfp-MapZ together with that of HlpA-mKate2, a chromosomal marker, in cells affected in chromosome segregation (Fig. 2E and Extended Data Fig. 3). Chromosome segregation defects were either induced by a treatment with the DNA-damaging agent mitomycin C^20^, or by the deletion of the gene coding for RocS, the protein that is required for proper segregation of *OriC* in the two daughter cells^19^. After treatment with a sub-lethal dose of MMC (7.5 ng/ml) or in the Δ*rocS* mutant, respectively 55±15% and 43±3% of cells harbor a chromosome that was either not or only partially segregated (Fig. 2E and Extended Data Fig. 3). However, the localization of FtsZ-mGfp, FtsA-mGfp or mGfp-MapZ at the cell equators were not affected in anucleate cells (Fig. 2E and Extended Data Fig. 3). Accordingly, the cell shape in both conditions remains largely unaffected with only 8±6% of misshaped cells. Time-lapse imaging of FtsZ-mGfp and HlpA-mKate2 in cells treated with MMC further showed that FtsZ-mGfp localize at the division site even in the absence of properly segregated DNA (Figure 2F and Video 2). Together, these results show that *OriC* or the nucleoid are not involved in the initial placement of the division site.

### The localization of MapZ at the cell equators is modified in absence of DacA or DacB

The localization of MapZ at the cell equators site is driven by the direct interaction between its extracellular domain and the PG^9,12^. To uncover the molecular determinants responsible for the localization of the extracellular domain of MapZ at the cell equators, we focused on two PG hydrolases, the carboxypeptidases DacA and DacB, whose inactivation has been associated with misplaced division septa^21–25^. As previously observed, the growth and the morphology of the Δ*dacA* and Δ*dacB* mutants were strongly affected, exhibiting a highly variable cell length and width, with a significant proportion of cells exhibiting asymmetrical shapes (Fig. 3A and Extended Data Fig. 4). As expected, the localization of FtsA-mKate was strongly altered in a *ΔdacA* or a *ΔdacB* mutant (Fig. 3A and Extended Data Fig. 5A), confirming that DacA and DacB are therefore required for the correct localization of the divisome at the cell equators^22,24^. These phenotypes were completely abolished upon ectopic expression of *dacA* or *dacB* (Extended Data Fig. 5B) and were also observed in the corresponding catalytic mutants (dacA_S56A_ and dacB_E204A_) (Extended Data Fig. 5A&C). This confirm that the activities of DacA and DacB are crucial for the correct placement of the division site in *S. pneumoniae*.

**Figure 3:**
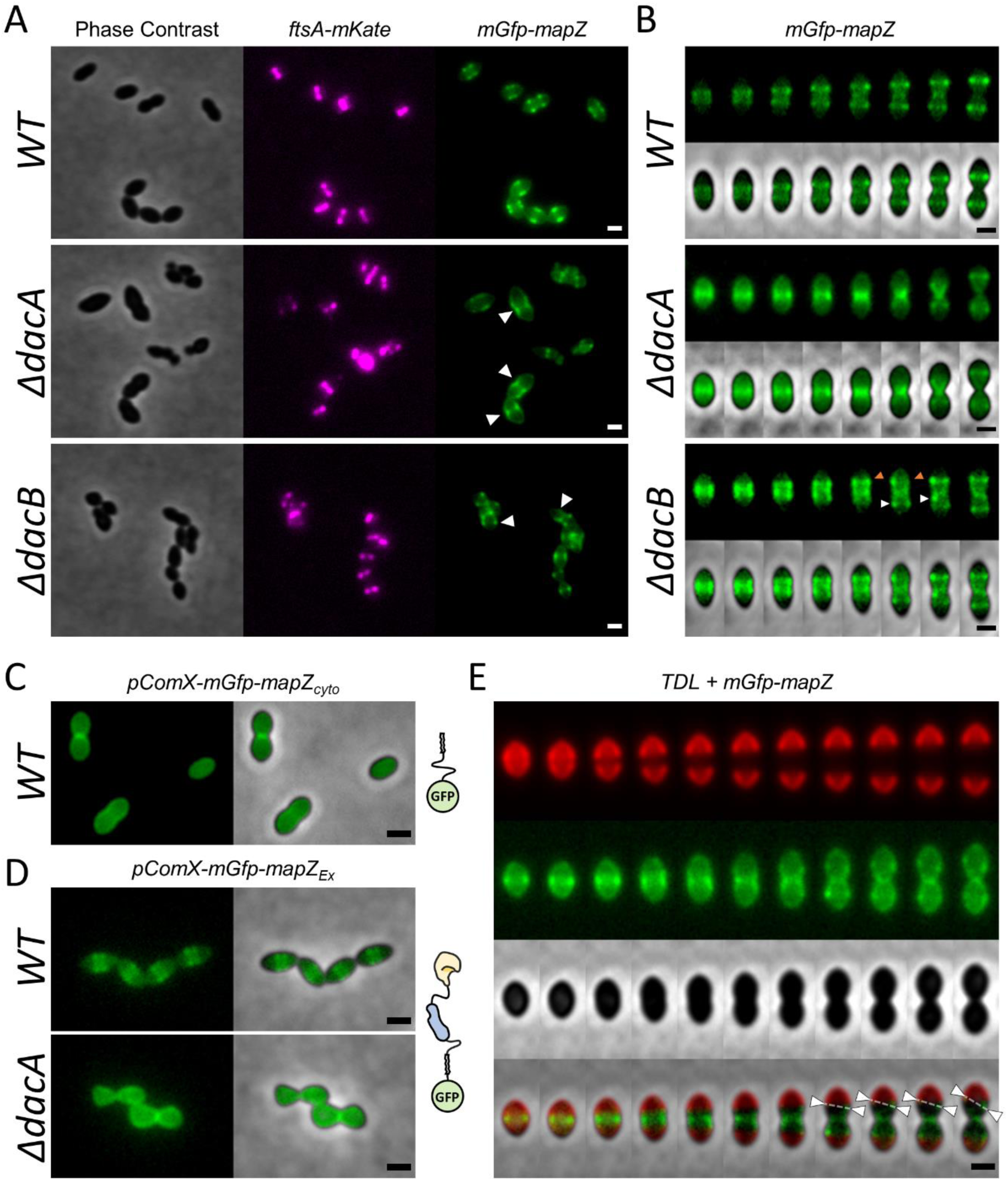
The localization of MapZ at the cell equators is strongly affected in the absence of the carboxypeptidase activities of DacA and/or DacB. (A) Representative images showing the localization of FtsA-mKate and mGfp-MapZ in WT, ΔdacA and ΔdacB mutant cells. The white triangles highlight the abnormal mGfp-MapZ localizations. The phase contrast, the mKate and the mGfp channel images are shown. The scale bar represents 0.5 µm. (B) Representative montages of images showing the localization of mGfp-MapZ as the cell elongates at the single cell level by time-lapse microscopy in WT, *ΔdacA* and *ΔdacB* mutant cells. The white triangles highlight a MapZ ring whose localization is erratic. The orange triangles highlight a MapZ ring whose formation is inconstant. Pictures were taken every 2 min. The mGfp channel and the merge between the mGfp channel and the phase contrast images are shown. The scale bar represents 0.5 µm. (C) Representative images showing the localization of the truncated version of MapZ with only the cytoplasmic domain (mGfp-MapZ_cyto_) expressed under the control of an inducible promoter in WT cells. (D) Representative images showing the localization of the truncated version of MapZ with only the extracellular domain (mGfp-MapZ_Ex_) expressed under the control of an inducible promoter in WT and ΔdacA mutant cells. The mGfp channel and the merge between the mGfp channel and the phase contrast images are shown. The scale bar represents 0.5 µm. A schematic of the topology is shown for each version of MapZ where the first part of the extracellular domain (MapZ_Ex1_) is colored in light blue and the second part of the extracellular domain (MapZ_Ex2_) is colored in yellow. (E) A representative montage of images showing the localization of mGfp-MapZ and the interface between the new and the old poles (aka cell equators) as the cell elongates at the single cell level by time-lapse microscopy. The cell equators can be visualized using a long pulse of the fluorescent D-amino acid (FDAA) followed by a chased experiment performed under the microscope. Pictures were taken every 2 min. The TDL-channel, the mGfp-channel, the phase contrast and the merged images are shown. The dotted lines and the white triangles highlight a MapZ ring that is no longer positioned at the interface between the new and the old poles in a *ΔdacB* mutant. The scale bar represents 1 µm.

Interestingly, the localization and the dynamic of mGfp-MapZ were also significantly affected in the *ΔdacA* and the *ΔdacB* mutants (Fig. 3A&B). Upon deletion of *dacA,* mGfp-MapZ mostly localized at the active division site throughout the cell cycle, while it is usually observed at the cell equators in wild-type cells (Fig. 3A&B). Specifically, mGFP-MapZ re-localized to the future site of division only during the later stages of the cell cycle of Δ*dacA* cells, mirroring the dynamics of FtsZ and more broadly, the divisome^26^ (Fig. 3B). Since, the cytoplasmic domain of MapZ has been shown to interact with FtsZ through its cytoplasmic domain^9,12^, we looked at the sub-cellular localization of truncated variants of MapZ with only the transmembrane domain and either the cytoplasmic (mGfp-MapZ_cyto_) or the extracellular (mGfp-MapZ_Ex_) domain. These variants (as well as the others presented below) were stable and did not affect the cell morphology (Extended Data Fig. 6). Accordingly, the MapZ variant containing only the cytoplasmic domain predominantly localized at the active division site in WT cells (Fig. 3C). In contrast, the variant with only the extracellular domain localized at the cell equators of WT cells but became delocalized throughout the cell periphery in Δ*dacA* mutant cells (Fig. 3D). Taken together, these findings confirm that the cytoplasmic domain of MapZ drives the aberrant localization of MapZ at the active division site in the absence of DacA, likely through its interaction with FtsZ. More importantly, these findings show that MapZ behaves like a variant lacking the extracellular domain in the absence of DacA, suggesting that this domain is no longer able to specifically bind to the cell equators of *dacA* mutant cells.

The localization of mGfp-MapZ was also affected in the Δ*dacB* mutant (Fig. 3A&B). However, in contrast to the *ΔdacA* mutant, mGFP-MapZ did not localize at the active division site in the absence of DacB. Instead, MapZ formed rings that were no longer positioned at the cell equators (Fig. 3A-B). In addition, while MapZ rings remain at the cell equators as wild-type cells elongate^9,10^ (Fig. 3B), they were no longer perpendicular to the cell’s longitudinal axis and continuously shifted in position and/or orientation in the absence of DacB, becoming stationary only during the later stages of the cell cycle (Extended Data Fig. 5D and Video 3). By tracking mGFP-MapZ localization alongside the interface between the old and new poles using FDAA, we confirmed that these MapZ rings were no longer positioned at the cell equators in the Δ*dacB* mutant (Fig. 3E). Finally, when we looked at the localization of mGfp-MapZ and FtsA-mKate2 in the Δ*dacB* mutant, we observed that they co-localized only in the later stages of the cell cycle (Extended Data Fig. 5D). Interestingly when they both co-localized, they exhibited similar erratic movements. This suggested that, despite the aberrant localization of MapZ in the Δ*dacB* mutant, the placement of the divisome is to some extent supported by MapZ (Extended Data Fig. 5D). These results suggest that in the absence of DacB, the localization of the extracellular domain of MapZ is no longer restricted to the cell equators. Therefore, the carboxypeptidase activities of both DacA and DacB are both necessary for positioning the extracellular domain of MapZ at the cell equators.

### Cell equators are specifically enriched in tetrapeptides

DacA removes the last D-Ala residue at the fifth position of the stem-peptide to produce a tetrapeptide which can be subsequently processed by DacB to generate a tripeptide^21,22,24,27^. Since the localization of MapZ is aberrant in the absence of DacB and completely disrupted in the absence DacA, we hypothesized that the binding of the extracellular domain of MapZ might require the presence of tetrapeptides at the cell equators (discussed below). The extracellular domain of MapZ displays a bi-modular structure composed of two subdomains, MapZ_Ex1_ and MapZ_Ex2_, separated by a flexible linker^9,12^. While the MapZ_Ex2_ sub-domain mediates the interaction with the PG, the MapZ_Ex1_ sub-domain acts as a pedestal, positioning MapZ_Ex2_ correctly in space and/or orientation relative to its ligand^9,12^. Accordingly, when expressed in wild-type cells, a truncated variant of MapZ lacking both the cytoplasmic domain and the MapZ_Ex2_ sub-domain (mGfp-mapZ_Δcyto_Δ_Ex2_) localized throughout the cell periphery (Fig. 4A). We reasoned that substituting the MapZ_Ex2_ sub-domain for a domain known to interact with tetrapeptides should allow to recover a localization at the cell equators. To test this hypothesis, we engineered a chimeric variant of MapZ lacking the cytoplasmic domain in which the MapZ_Ex2_ sub-domain was replaced by the inactive catalytic domain of DacB (mGfp-mapZ_Δcyto-Ex2::_DacB_E204A_). When expressed in WT cells, this chimeric version of MapZ harbored the same subcellular localization and dynamic than MapZ, *i.e.* a single ring located at the division site in the nascent cell that eventually split into 2 rings that migrated with the cell equators as the cell elongates (Fig. 4B&C). These results strongly suggest that tetrapeptides are specifically recognized by the MapZ_Ex2_ sub-domain of MapZ at the cell equators.

**Figure 4:**
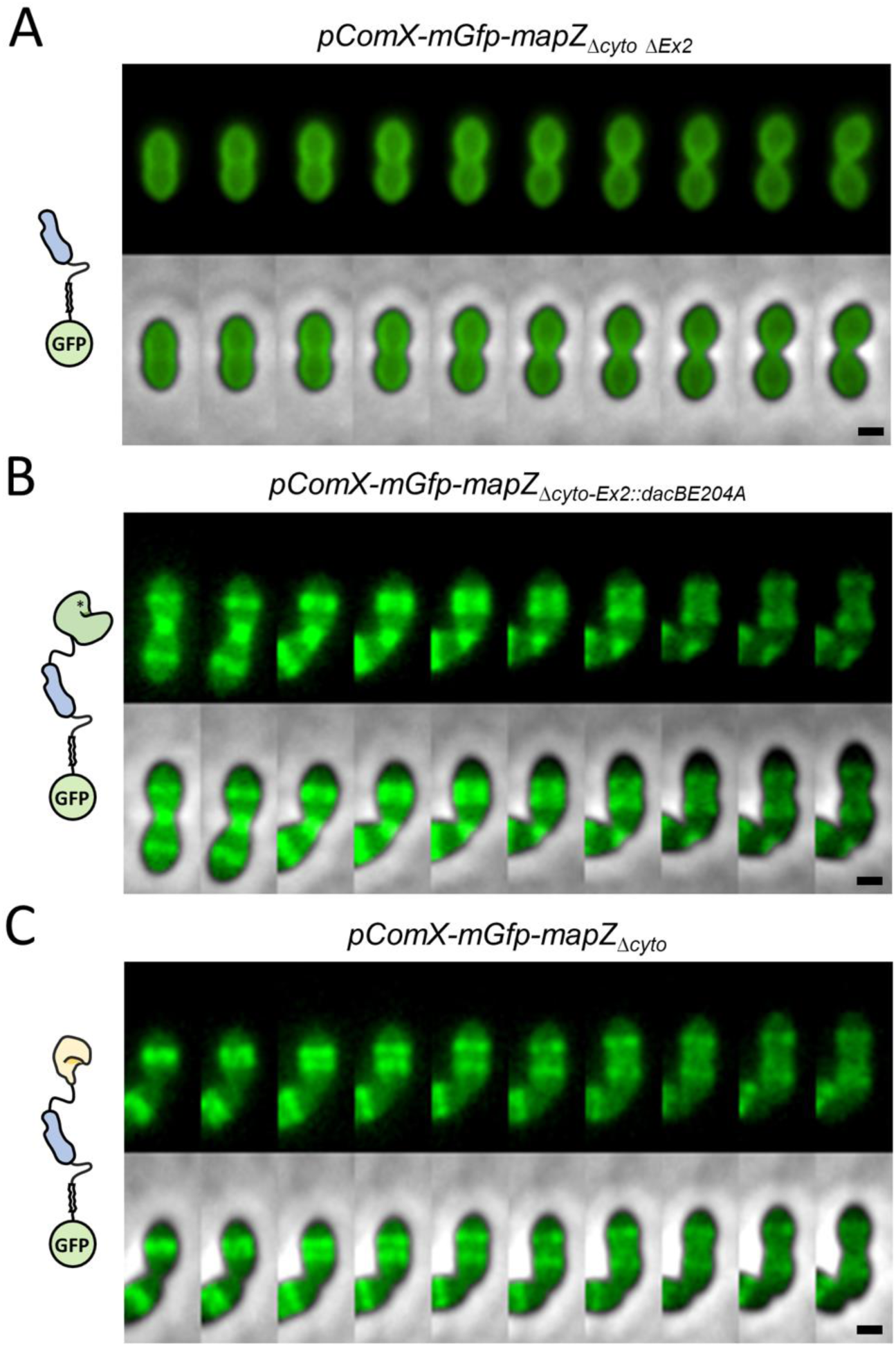
Cell equators must be specifically enriched in tetra-peptide. (A-C) Representative montage of images showing the localization and the dynamic of truncated variants of MapZ as the cell elongates at the single cell level by time-lapse microscopy. A variant lacking both the cytoplasmic domain and the MapZ_Ex2_ sub-domain (mGfp-mapZ_Δcyto_Δ_Ex2_) is shown in (A). The chimeric version of MapZ lacking only the cytoplasmic domain and where the Map_Ex2_ domain was replaced by the mutated catalytic domain of DacB is shown in (B). A variant lacking only the cytoplasmic domain is shown in (C). A schematic of the topology is shown for each version of MapZ where the first part of the extracellular domain (MapZ_Ex1_) is colored in light blue, the second part of the extracellular domain (MapZ_Ex2_) is colored in yellow and the catalytic domain of DacB is colored in green. The mGfp channel and the merge between the mGfp channel and the phase contrast images are shown. The scale bar represents 0.5 µm.

### DacA and DacB are sequentially recruited at the division site

As mentioned previously, tetrapeptides are mostly produced by DacA and serve as substrate for DacB to generate tripeptides^21,22,24,25,27^. The activity and/or localization of DacA and DacB must be precisely coordinated to generate tetrapeptides at the cell equators. Previous immunofluorescence microscopy studies indicated that DacA and DacB are distributed across the cell surface^22,24^. However, their dynamics throughout the cell cycle remained elusive. To gain more insight about the dynamic of DacA and DacB, we constructed corresponding fluorescent protein fusions and looked at their respective localization.

DacA is membrane-associated through an N-terminal amphipathic helix, followed by a globular domain that likely acts as a pedestal to position a second domain containing the active site^23^. Our attempts to generate N-or C-terminal sfGFP fusions showed that they were not functional. We therefore engineered another fusion, in which the sfGFP was inserted between the amphiphilic helix and the pedestal domain of DacA (DacA-sfGFP-H) (Extended Data Fig. 7A). On the other hand, DacB is a lipoprotein divided into an N-terminal disordered region and a C-terminal catalytic domain. Likewise, the N- and C-terminus fusions were not functional. We therefore inserted the sfGFP between the lipoprotein signal peptide and the disordered region of DacB (sfGFP-DacB) (Extended Data Fig. 7D). Both DacA and DacB fusions were expressed at the native chromosomal locus and were fully functional ensuring wild-type growth kinetics and normal cell morphology (Fig. 5A-B and Extended Data Fig. 7). To determine the dynamic of DacA and DacB throughout the cell cycle, we monitored the localization of DacA-sfGFP-H and sfGFP-DacB at the single cell level (Fig. 5A-B). Their averaged sub-cellular localization was monitored for four classes of cell length as a proxy for their progression through the cell cycle. DacA-sfGFP-H mostly localized at the division site throughout the cell cycle, forming a band that gradually increased in intensity as the septum closed. At the initial stage of the cell cycle, when cells had completed division and may not be yet separated, DacA-sfGFP-H was absent from the division site and was observed mostly at the equator of the daughter cells (Fig. 5A). By comparison, sfGFP-DacB was observed primarily as faint dots at one or both poles at the initial stage of the cell cycle and as a band at the division site that gradually increased in intensity as the septum closed (Fig. 5B). Consequently, when DacA-sfGFP-H was observed at the active division site in the early stage of the cell cycle, sfGFP-DacB was not present and appeared primarily as dots at one pole, which corresponds to the division septum of the previous cell cycle. This localization pattern suggests thus that DacB is recruited to the division site subsequent to DacA.

**Figure 5:**
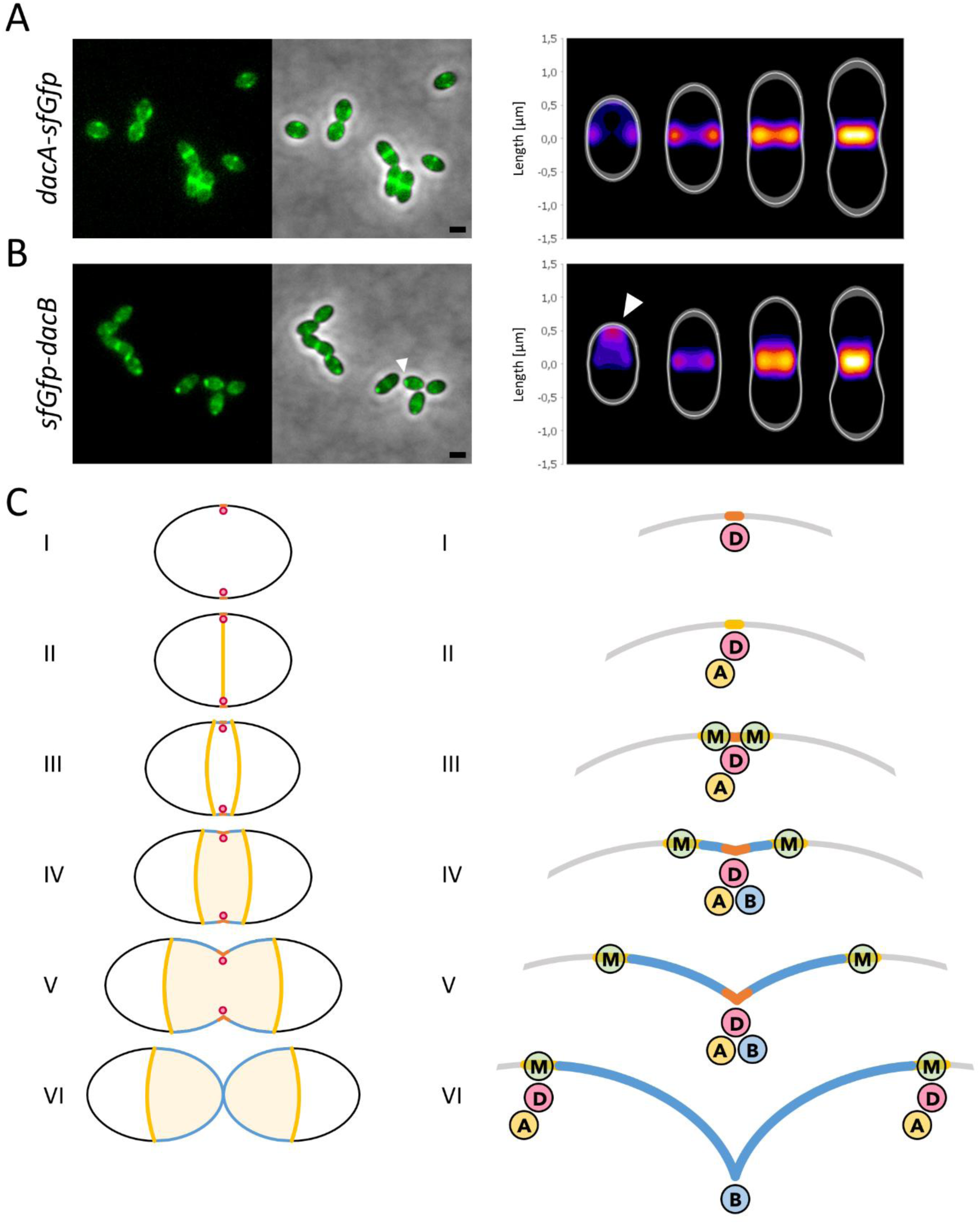
DacA and DacB are sequentially recruited at the division site. (A-B) Representative images showing the localization of dacA-sfGfp (A) and sfGfp-dacB (B) in WT cells. The mGfp channel and the merged between the mGfp channel and the phase contrast are shown. The averaged sub-cellular localization was monitored for 4 classes of cell length as a proxy for their progression through the cell cycle and shown as heatmaps for dacA-sfGfp (A, n=1800) and sfGfp-dacB (B, n=2700). The color saturation represents the density of the localizations using a Fire LUT. The scale bar represents 1 µm. (C) A model showing the formation of a specific PG composition at the cell equators and produced only during the early stages of cell division. As division begins (I), the divisome (pink circle) localizes within a ring-shaped region at the cell equators of the newly formed daughter cells and starts producing new PG (Orange). In the presence of DacA (yellow circle) and in the absence of DacB (blue circle), most pentapeptides are either cross-linked by PBPs or trimmed by DacA, resulting in a PG ring enriched with tetrapeptides (Yellow). This equatorial PG ring, enriched with tetrapeptides, is specifically recognized by MapZ marking the cell equators of the future daughter cells throughout the cell cycle (III). As new material is continuously added, this initial PG is gradually pushed outward from the division site, placing it beyond the reach of DacB by the time it is eventually recruited to the division site (IV). In the presence of DacB, most tetrapeptides are trimmed, resulting in a PG enriched with tripeptides (blue). Finally, as septum formation completes and cell separation occurs, PG synthesis proteins relocate to the MapZ rings at the equators of the newly formed daughter cells to initiate a new cycle.

## Discussion

Misplacement of the division site can lead to asymmetrical division, resulting in cells that lack essential cellular components, which can impair survival and cell function^28,29^. Here we demonstrated that the placement of the division site in the ovoid shaped bacteria *S. pneumoniae* does not occur at mid-cell but rather at the cell equator, the widest part of the cell body. We confirmed that only MapZ localizes throughout the cell cycle at the cell equators where it will eventually recruit the divisome to initiate a new round of cell division. On the other hand, the localization of MapZ and the divisome at the cell equators were not affected in the absence of properly segregated DNA suggesting that OriC or the nucleoid are not involved in division site selection as suggested before^11^. We observed that MapZ localization is significantly altered in the absence of the two PG hydrolases DacA and/or DacB. Furthermore, we provided evidence that these two enzymes create a distinct signature at the future division sites, likely involving tetrapeptides, which are crucial for positioning MapZ at the cell equators.

PG is composed of glycan strands made of repeating units of N-acetylglucosamine (GlcNAc) and N-acetylmuramic acid (MurNAc). Short peptides comprising up to five amino acids (L-alanyl-γ-D-glutamyl-L-lysyl-D-alanyl-D-alanine) are attached to the MurNAc residues. These peptides are cross-linked between the D-alanine at the fourth position of a stem peptide (donor peptide) with the L-lysine at the third position in one adjacent stem peptide (acceptor peptide), thereby forming a robust, mesh-like polymeric matrix^30^. The synthesis of PG involves penicillin-binding proteins (PBPs), and SEDS (shape, elongation, division, and sporulation) proteins^31^. In many bacteria, these proteins work in concert with PG hydrolases, such as DacA and DacB, to remodel the peptidoglycan and regulate its crosslinking, ensuring the formation of cells with proper shape and size ^21,22,24,25^. DacA and DacB are two carboxypeptidases that act in a sequential manner on the stem peptides. DacA trims the terminal D-alanine residues from the pentapeptide leading to a tetrapeptide, which is further processed by DacB to form a tripeptide^21,22,24,25,27^. Because only pentapeptides can serve as donor peptides and tripeptides seems to be preferred as acceptor peptides, carboxypeptidases are believed to regulate PBP transpeptidase activity by controlling substrate availability^22^. Previous studies indicate that the primary monomeric and cross-linked PG peptide species isolated from WT cells are tripeptides, suggesting that most of the pentapeptides are either crosslinked by the PBPs or trimmed by DacA, and then processed by DacB^21,22,24^. Accordingly, pentapeptides accumulate in the PG of a *ΔdacA* mutant, while tetrapeptides are notably absent^21,22,24^. Interestingly, MapZ no longer localized at the cell equators in the *ΔdacA* mutant (Fig. 3A&B), and behaves like a variant lacking the extracellular domain (Fig. 3C). Consequently, in the absence of DacA, where tetrapeptides are nearly absent, the extracellular domain can no longer bind to the cell equators. Conversely, in a *ΔdacB* mutant, where tetrapeptides accumulate^21,22,24^, MapZ rings were unstable, frequently shifting in position and/or orientation as the cell elongated (Fig. 3B&D). Under these conditions, the unstable localization of MapZ may result from an excess of tetrapeptides that could be no longer confined to the cell equators. Finally, a chimeric version of MapZ, in which the extracellular domain MapZ_Ex2_ was replaced by a domain known to interact with tetrapeptides, mimicked the equatorial localization of MapZ (Fig. 4D). Altogether these results strongly suggest that tetrapeptides enriched at the cell equators are specifically recognized by the extracellular domain of MapZ and are essential for its positioning.

The specific binding of MapZ to tetrapeptides at the cell equators raises the question of how tetrapeptides become specifically enriched at this subcellular location. The answer most likely lies in the unique mechanism of cell constriction and elongation exhibited by the pneumococcus. Unlike rod-shaped bacteria, which incorporate new peptidoglycan either at the division septum or along their lateral walls, *S. pneumoniae* exclusively incorporates new material at the division site^17^. This process supports both cell elongation and constriction, ultimately forming a new cell half at each division cycle^32,33^. Short pulses with fluorescent D-amino acid (FDAA) have shown that the new PG is incorporated circumferentially, with the oldest material gradually displaced outward from the division site, due to the continuous addition of new material^32,33^. As a result, the cell equator always lies at the junction between the newly synthetized cell half and the old cell half inherited from the mother cell. This mode of growth implies that the ring of PG at the cell equators is synthesized at the very beginning of the cell division process. By using functional fluorescent fusions in living cells, we demonstrated that DacA and DacB both localize mostly at the active division site (Fig. 5), like the main components of the divisome such as the PBPs^26,34^. However, our analysis revealed a slight delay in their recruitment to the future site of division (Fig. 5). Notably, DacA is recruited to the cell equators in the early stages of division, as peptidoglycan synthesis initiates whereas DacB displays a dynamic characteristic of a late-division protein^26^. Indeed, in the early stages of division, DacB was still observed at the old division site forming a faint dot at one pole while DacA and most of the divisome localized at the cell equators of the new daughter cells. This suggests that DacB is recruited to the division site after DacA, thereby preserving tetrapeptides in the early stages of division. In this model, as division begins, PG synthesis initially take place within a ring-shaped region at the cell equators of the newly formed daughter cells (Fig. 5C). In the absence of DacB, most pentapeptides are either cross-linked by PBPs or trimmed by DacA, resulting in a PG ring enriched with tetrapeptides. As new material is continuously added, this initial PG is gradually pushed outward from the division site, placing it beyond the reach of DacB by the time it is eventually recruited to the division site. This equatorial PG ring, enriched with tetrapeptides, is specifically recognized by MapZ marking the cell equators of the future daughter cells throughout the cell cycle to eventually recruit FtsZ. Finally, as septum formation completes and cell separation occurs, PG synthesis proteins relocate to the FtsZ rings at the equators of the newly formed daughter cells to initiate a new cycle. Altogether, this model shows that the sequential recruitment of DacA and DacB to the division site during early PG synthesis could drive the formation of local PG composition at the cell equators. In conclusion, our findings provide valuable insights toward the ultimate characterization of the structural and functional characteristics of the specific signature recognized by MapZ.

## Materials and Methods

### Strains and growth conditions

Strains used in this study are listed in the Table S1. *Streptococcus pneumoniae R800* and derivatives were cultivated at 37°C in C+Y medium at pH 7.4. Cell growth was monitored automatically in JASCO V-630-BIO-spectrophotometer by optical density (OD). Cells were grown in C+Y medium at pH 7.4 until they reached early log phase (OD_550nm_=0.1-0.2) and then normalized at OD_550nm_=0.001 in the corresponding medium. Optical density was read automatically at 550 nm every 10 min for 1000 min.

### Allelic replacement mutagenesis

Gene modifications in *S. pneumoniae* were achieved by homologous recombination using the two-step procedure based on a bicistronic *kan-rpsL* cassette called Janus^35^ and constructed at their native chromosomal locus. Briefly, the Janus cassette is used to replace the gene of interest. After transformation and selection, this confers resistance to kanamycin and dominant streptomycin sensitivity (Kan^R^–Str^S^) in an initial WT *rpsL1* genetic background. Then, a DNA fragment flanked on each end by sequences homologous to the upstream and downstream regions of the gene of interest is used to transform and substitute the Kan^R^–Str^S^ in the Kan^R^ strains. Integration of the fragment is selected in final non-polar markerless mutant strains by streptomycin resistance. *S*. *pneumoniae* mutants were constructed by transformation of the WT strain or derivatives using pre-competent cells treated at 37°C during 30 min with the synthetic competence stimulating peptide 1 (CSP1) at the concentration of 100 ng/10^8^ cells to induce competence. Cells were then plated on THY-agar supplemented with 3% (vol/vol) defibrinated horse blood and incubated for 120 min at 37°C. Selection of transformants was then performed by adding a THY-agar overlay containing the appropriate antibiotic (streptomycin 200 μg/mL, kanamycin 250 μg/mL) and further incubated for 16 h at 37°C. The oligonucleotides used for all construction are listed in Table S2. Pneumococcal strains were verified by DNA sequencing to verify error-free PCR amplification.

### Microscopy techniques

Cells were grown until OD_550nm_ = 0.1 and spotted on pads made of 1% agarose in C+Y medium at 37°C. Slides were visualized with a Nikon Ti-E microscope fitted with an Orca-CMOS Flash4 V2 camera with a 100 × 1.45 objective. Images were collected using NIS-Elements (Nikon). Images were analyzed using the software ImageJ (http://rsb.info.nih.gov/ij/) and the plugin MicrobeJ^18^.

FtsZ-mGFP, mGFP-MapZ or FtsA-mKate rings were detected using the feature/strip detection option in MicrobeJ. This option combines spatial 1D filtering (Median Filter) and 1D local maxima algorithm to localize local maxima in the fluorescent profile extracted along the medial axis of each detected cell. Each local maximum was then fit to a single peak or a multi peak 1D Gaussian curve, to determine their amplitude, their FWHM (Full width at half Maximum) and their coordinates at the subpixel resolution. Local maxima were finally filtered based on the goodness of the fit and their amplitude. Their sub-cellular localizations along the longitudinal axis were automatically computed for each associated particle.

Diffraction-limited foci of ParB-mGFP were detected using the feature/spot detection option in MicrobeJ as previously described^19^. Briefly, this option combines spatial 2D filtering (Median Filter) and 2D local maxima algorithm to localize single fluorescent maxima in each detected cell. Each maximum was then fit to a single peak or a multi peak 2D Gaussian curve, to determine their amplitude, their FWHM and their coordinates at the subpixel resolution.

For FDAA staining, pneumococcal cells were grown until OD550nm = 0.1/0.2 and incubated for 3 h at 37°C in C+Y with 500 mM of TAMRA-D-lysine (TDL) or 7-hydroxycoumarincarbonylamino-D-alanine (HADA). Cultures were washed three times with 1 ml of cold C+Y and then resuspended in 50 μL cold C+Y before observation.

### Preparation of *S. pneumoniae* crude extracts and immunoblot analysis

Cultures of *S. pneumoniae* were grown in C+Y, pelleted and resuspended in Tris-HCl 10mM pH 8, EDTA 1mM. After cell disruption by sonication, crude extracts were analyzed by SDS-PAGE and electro transferred onto an immobilon-P membrane (Millipore). Primary antibodies were used at 1: 5000 (anti-GFP, Amsbio), in TBST-BSA 1% and 1: 250000 (anti-Enolase) in TBST-BSA 5%. The goat anti-rabbit secondary antibody HRP conjugate (Biorad) was used at 1: 5 000.

## Author Contributions

AD, and CG designed research; AD and CF performed experiments; AD, CF and CG analyzed data; AD and CG wrote the manuscript. All authors revised the manuscript.

### Acknowledgments

We thank Thomas Becker for expert technical assistance. Support for this work comes from the CNRS, the Université Lyon I, the foundation Bettencourt-Schuller to CG and the Agence Nationale de la Recherche (ANR-23-CE11-0029 and ANR-19-CE15-0011 to CG). We acknowledge the contribution of the SFR Biosciences (Université Claude Bernard Lyon 1, CNRS UAR3444, INSERM US8, ENS).

---

**Extended Data Figure 1:**
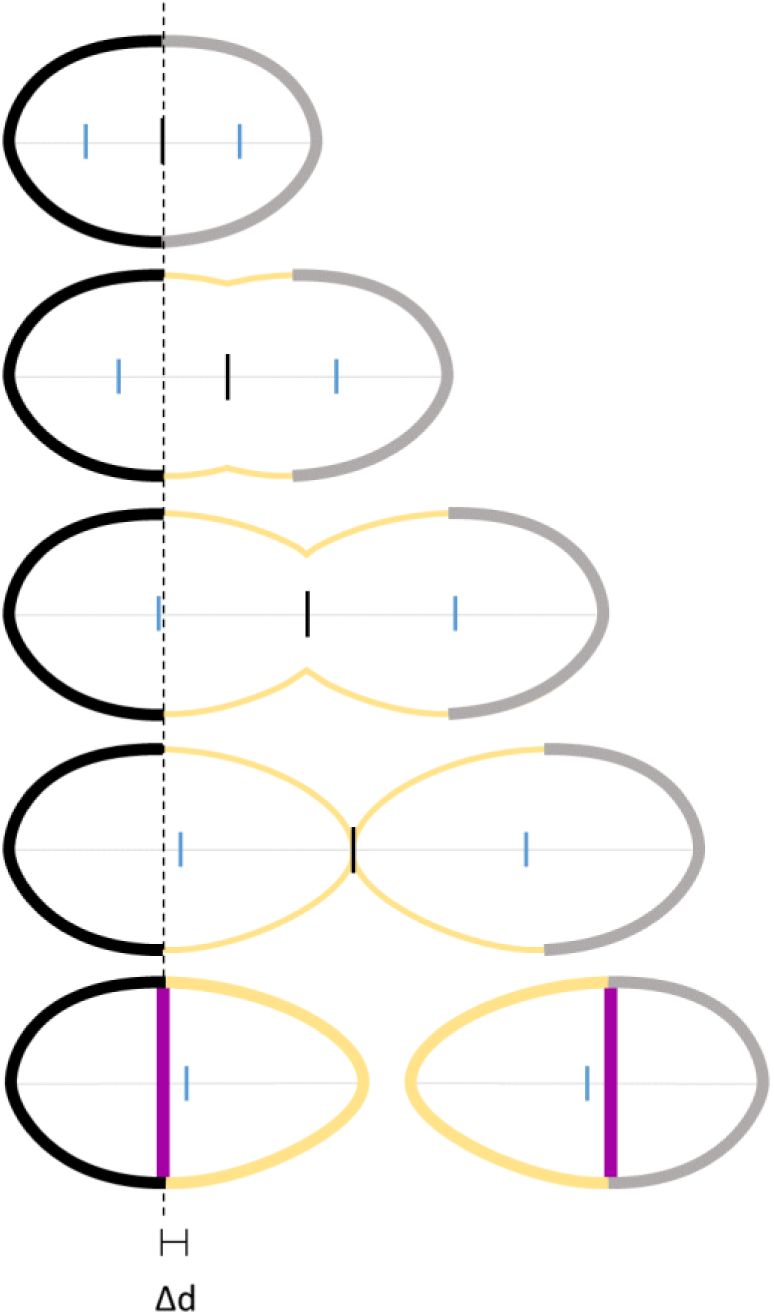
The formation of asymmetrical cells is driven by inherent regulatory variations between constriction and elongation across successive division cycles. *S. pneumoniae* incorporates new material (yellow) exclusively at the division site, forming a new cell pole at each division cycle. Consequently, the daughter cells always consist of a newly synthetized pole (yellow) and an old pole inherited from the mother cell (black or gray). The formation of asymmetrical cells is thus likely driven by inherent regulatory variations across successive division cycles. This also infers that the widest part of the cell, also known as the cell equators (black dotted line), always lies at the junction between the newly synthetized pole and the old pole inherited from the mother cell. Note that when the daughter cell is asymmetric, the cell equator is significantly offset relative to mid-cell (light blue dash).

**Extended Data Figure 2:**
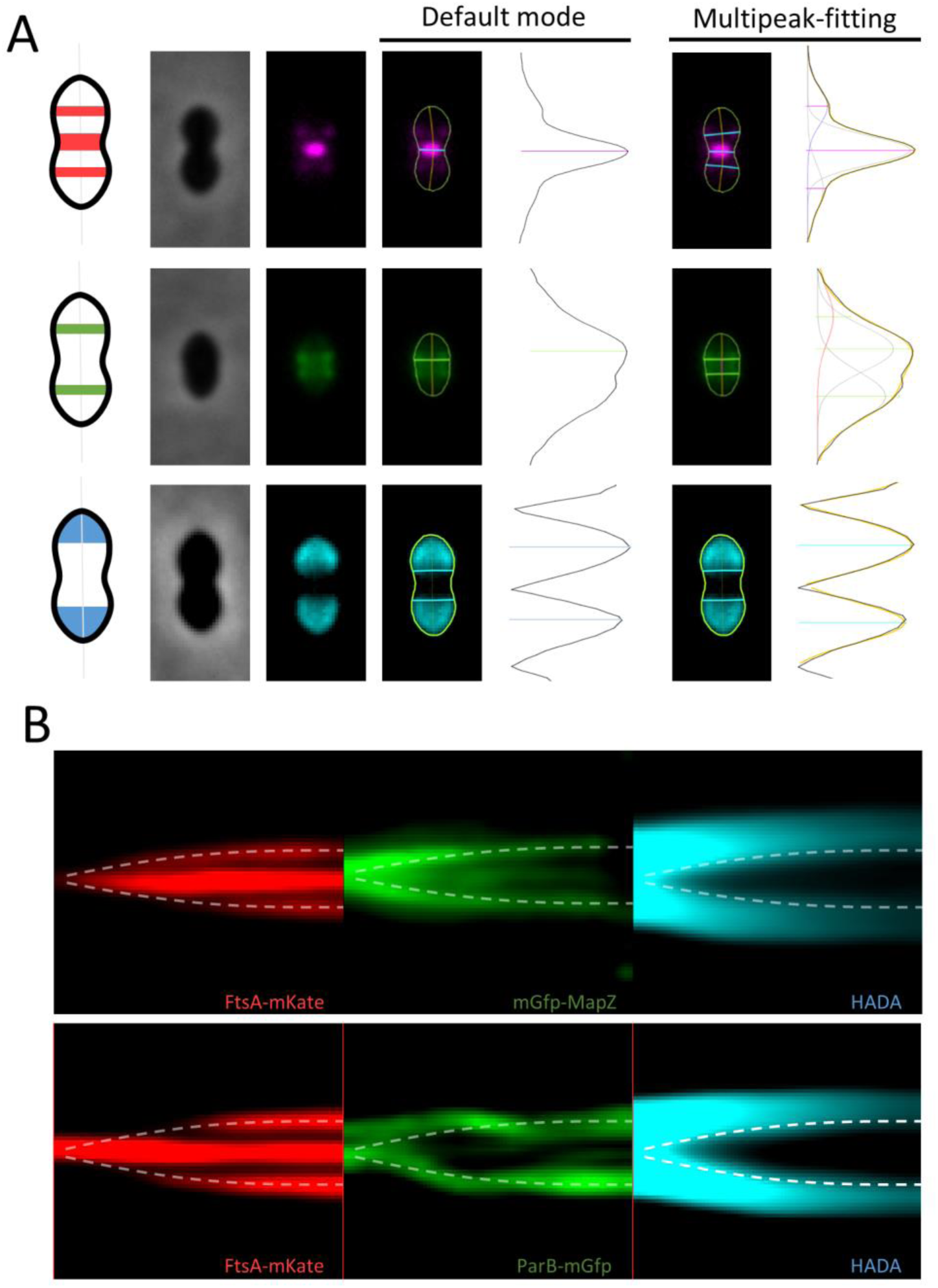
(A) Three representative images showing the detection of mKate2-FtsA, mGfp-MapZ rings and HADA staining by MicrobeJ using either the default mode (left side) or multipeak-fitting approach (right side). The phase contrast, mKate-channel, the gfp-channel or the HADA-channel images and the corresponding overlay and the longitudinal profiles retrieved from MicrobeJ are shown. (B) Representative kymographs showing the longitudinal dynamic of FtsA-mKate, mGfp-MapZ or ParB-mGfp, and the cell equators as the cell elongates at the single cell level by time-lapse microscopy. The white dash line represents the position of the cell equators deduced from the position of the interface between the old poles (HADA-labeled) and the new poles (unlabeled). The mKate, the mGfp and the HADA channel images are shown.

**Extended Data Figure 3:**
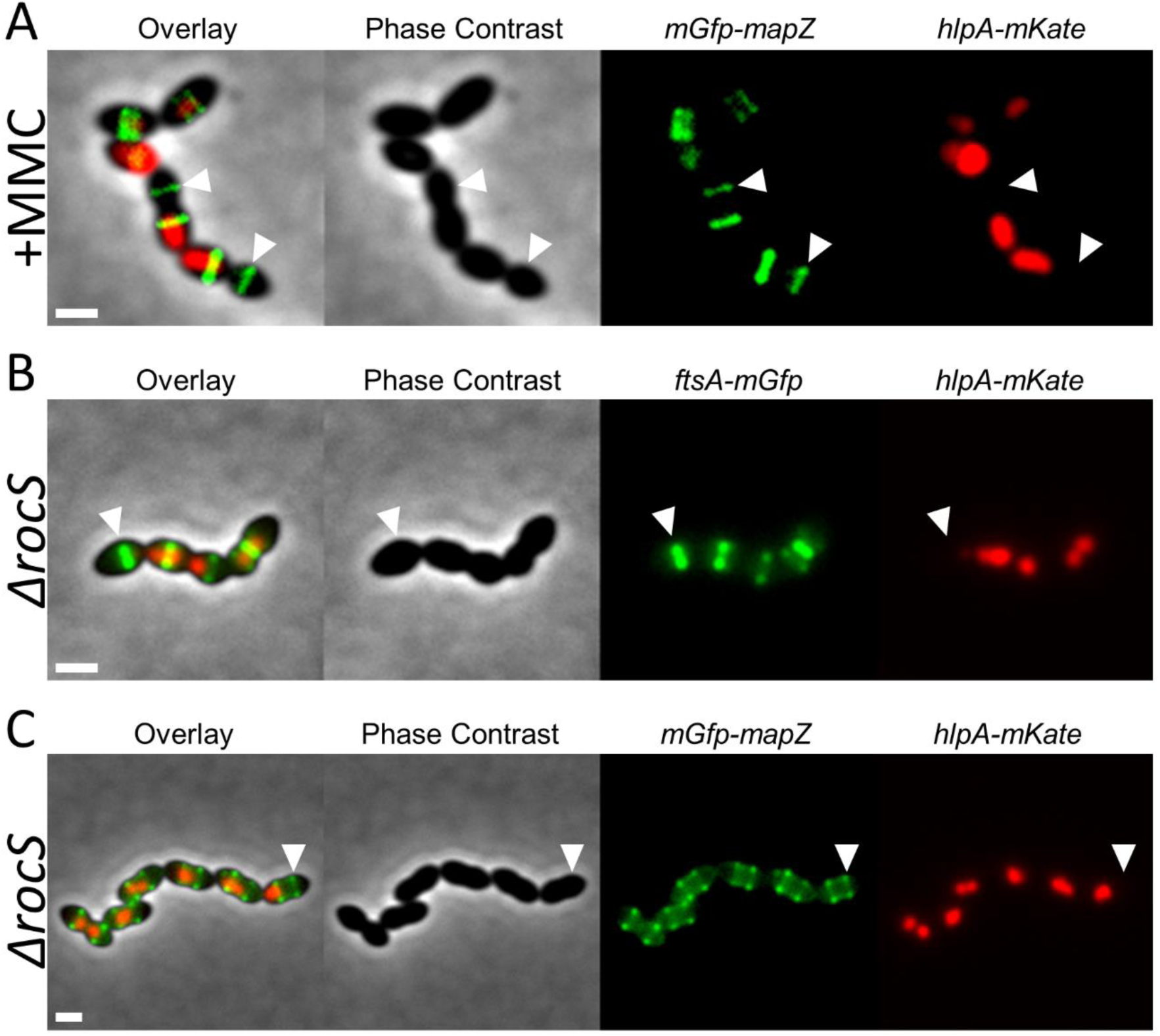
(A) Representative image showing the localization of mGfp-MapZ at the cell equators in cells whose chromosome (detected using an HlpA-mKate2 fusion) was either not or only partially segregated after treatment with a sub-lethal dose of MMC (7.5 ng/ml). The white triangles highlight the MapZ rings at the cell equators of cells where the nucleoid is not detectable. (B) Representative image showing the localization of FtsA-mGfp at the cell equators in cells whose chromosome (detected using an HlpA-mKate2 fusion) was either not or only partially segregated upon deletion of *rocS*. The white triangles highlight the FtsA rings at the cell equators of cells where the nucleoid is absent. (C) Representative image showing the localization of mGfp-MapZ at the cell equators in cells whose chromosome was either not or only partially segregated upon deletion of *rocS*. The white triangles highlight the MapZ rings at the cell equators of cells where the nucleoid is not detectable. (A-C) The merged of the phase contrast, the mKate-channel and the gfp-channel images, and the phase contrast images are shown. The scale bar represents 0.5 µm.

**Extended Data Figure 4:**
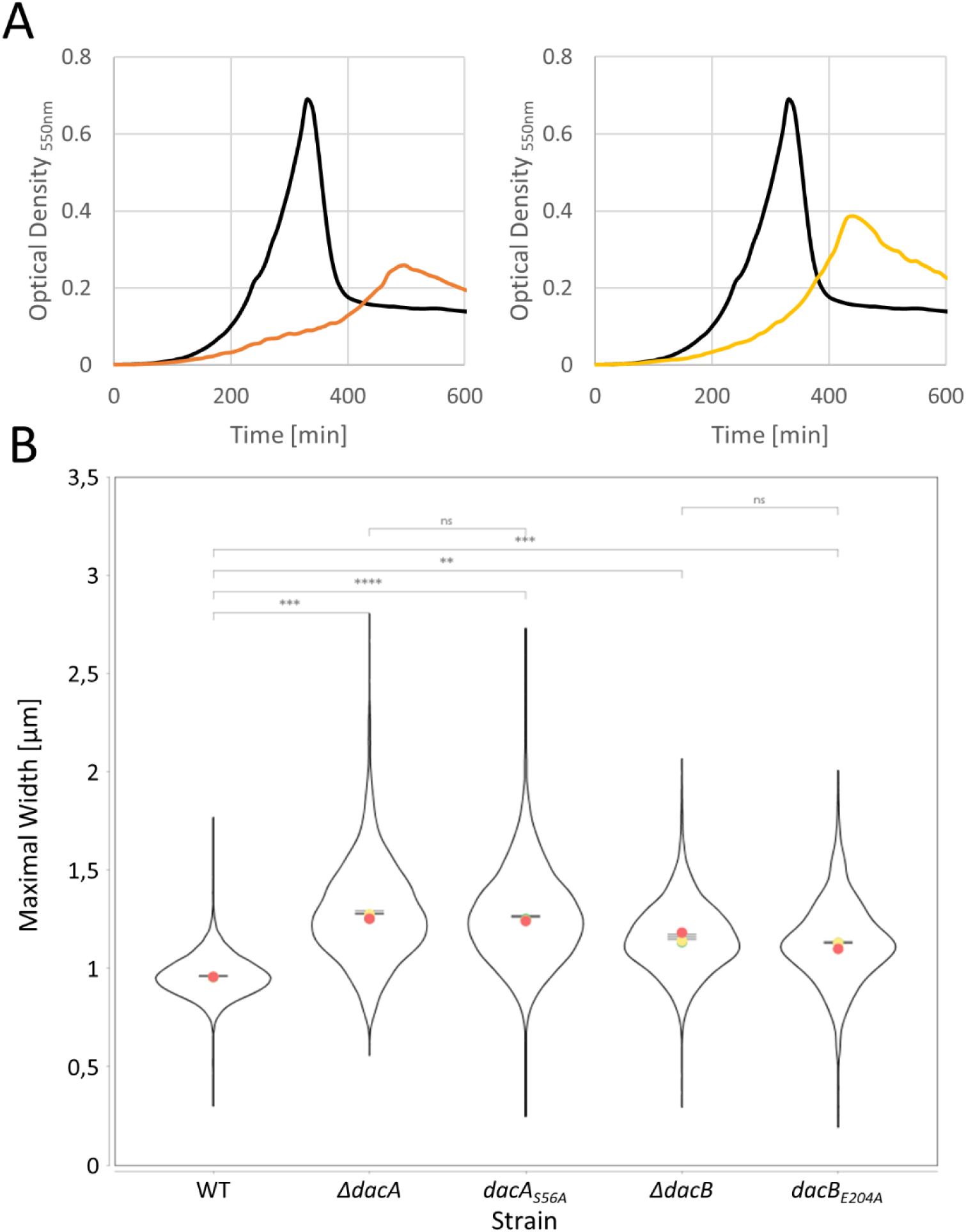
(A) Representative growth curves of the WT (solid line, black) and the *ΔdacA* mutant (solid line, orange) strain or the *ΔdacB* mutant (solid line, yellow) strain. (B) Super plots representing the distribution of the maximal width of WT cells or *ΔdacA*, *dacA_S56A_*, *ΔdacB or dacBE204A* mutant cells (n=25845, r=3, Unpaired T-test).

**Extended Data Figure 5:**
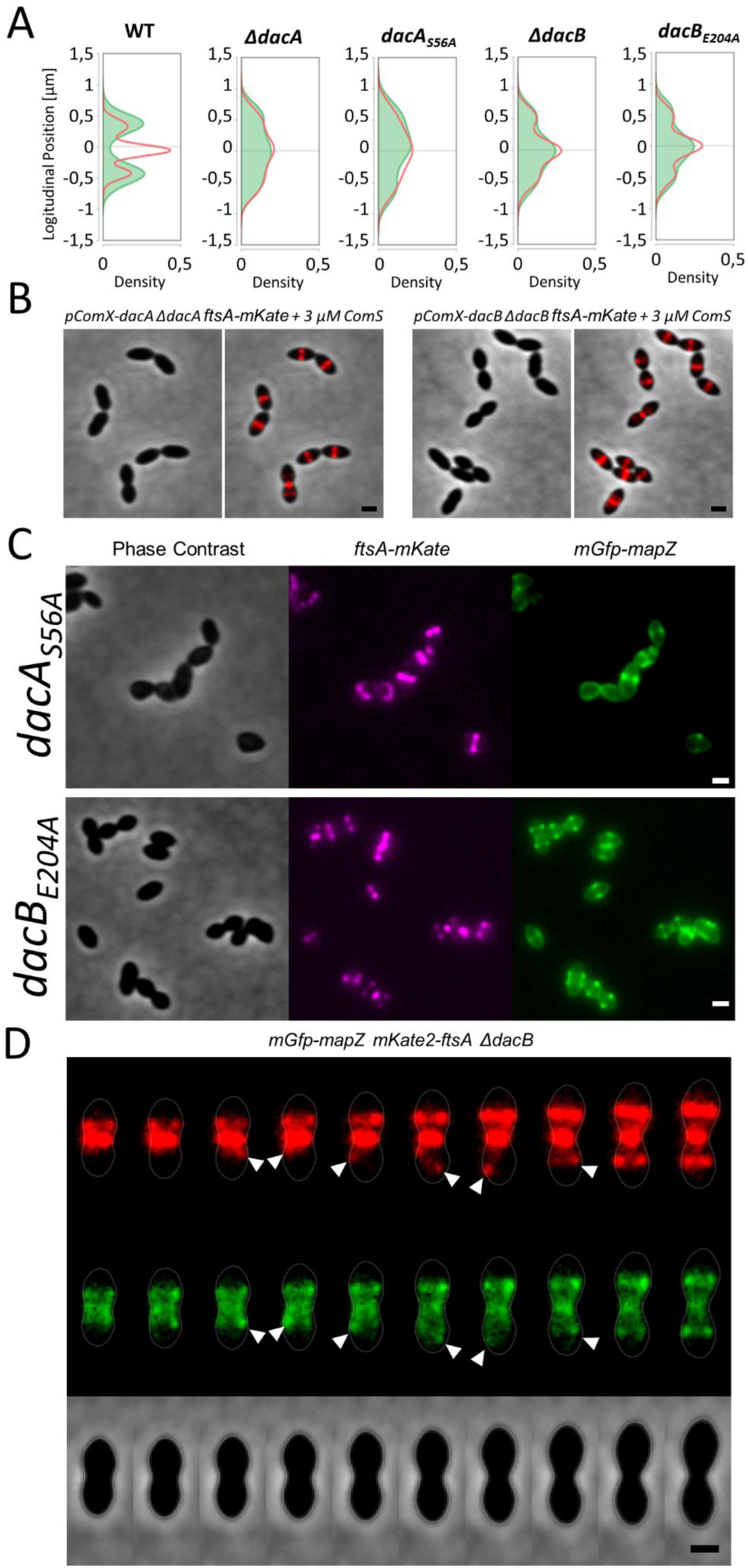
(A) The relative longitudinal position of FtsA-mKate (red line) and mGfp-MapZ rings (green line) shown as density plots for WT or *ΔdacA*, *dacA_S56A_*, *ΔdacB or dacBE204A* mutant cells with a cell length ranging from 1.8 µm to 2 µm. (B) Representative images showing the morphology and the localization of ftsA-mKate in *ΔdacA* (left panel) and *ΔdacB* (right panel) mutant cells upon complementation of *dacA* or *dacB* respectively. The phase contrast and the merged between the phase contrast and the mKate channel images are shown. The scale bar represents 0.5 µm. (C) Representative images showing the localization of FtsA-mKate and mGfp-MapZ in dacA_S56A_ and dacB_E204A_ mutant cells. The phase contrast, the mKate and the mGfp channel images are shown. The scale bar represents 0.5 µm. (D) A representative montage of images showing the localization and the dynamic of FtsA-mKate and mGfp-MapZ as the cell elongates at the single cell level by time-lapse microscopy in a *ΔdacB* mutant cell. Pictures were taken every 2 min. The mKate-channel, the mGfp-channel and the phase contrast are shown. The solid line represents the cell contour retrieved from MicrobeJ. The scale bar represents 1 µm.

**Extended Data Figure 6:**
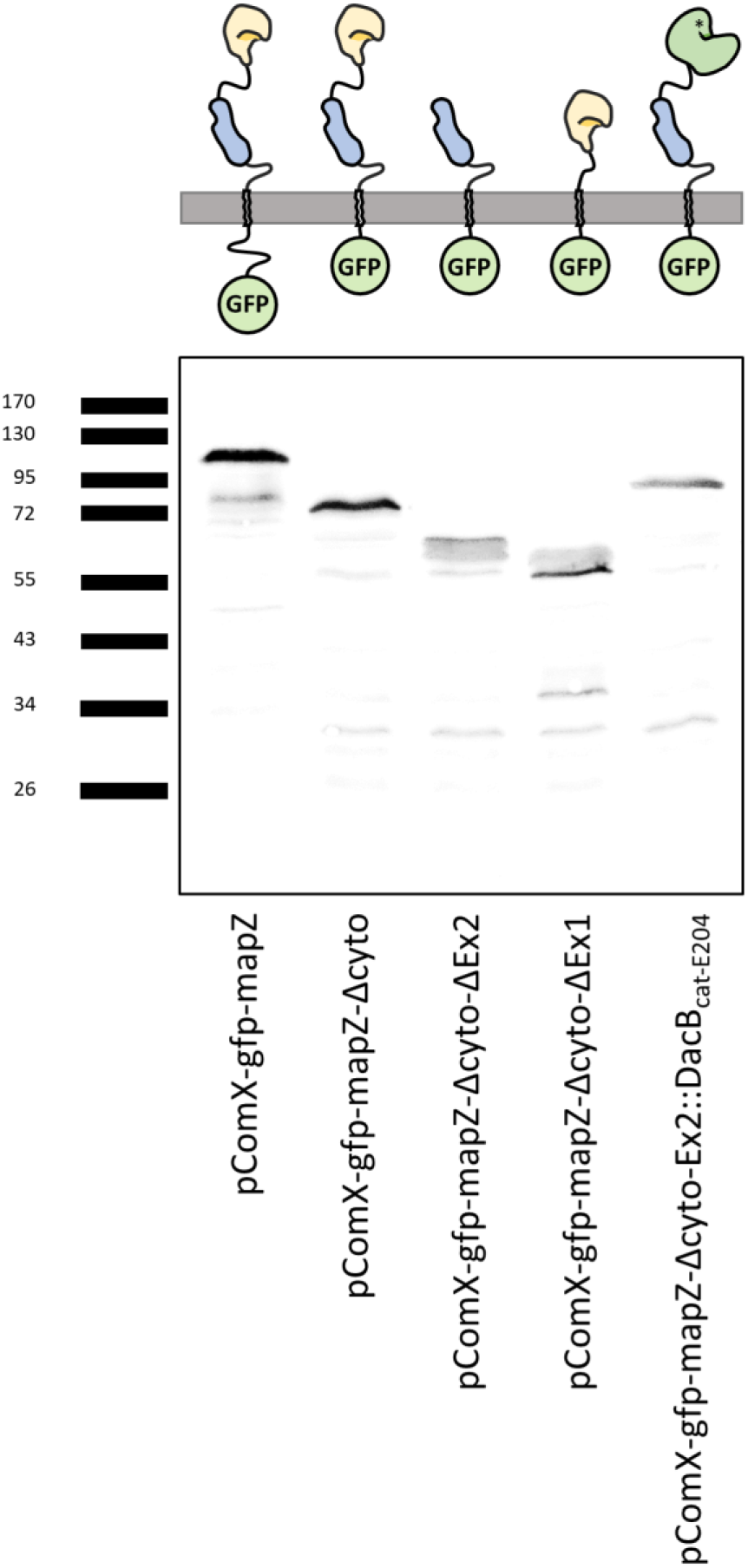
Western blot analysis of the different versions of MapZ, fused to the mGfp and expressed under the control of an inducible promoter in cells having an endogenous copy of *mapZ*. The fusions were detected with an anti-GFP antibody. A schematic of the topology is shown for each version of MapZ where the first part of the extracellular domain (MapZ_Ex1_) is colored in light blue, the second part of the extracellular domain (MapZ_Ex2_) is colored in yellow and the catalytic domain of DacB is colored in green.

**Extended Data Figure 7:**
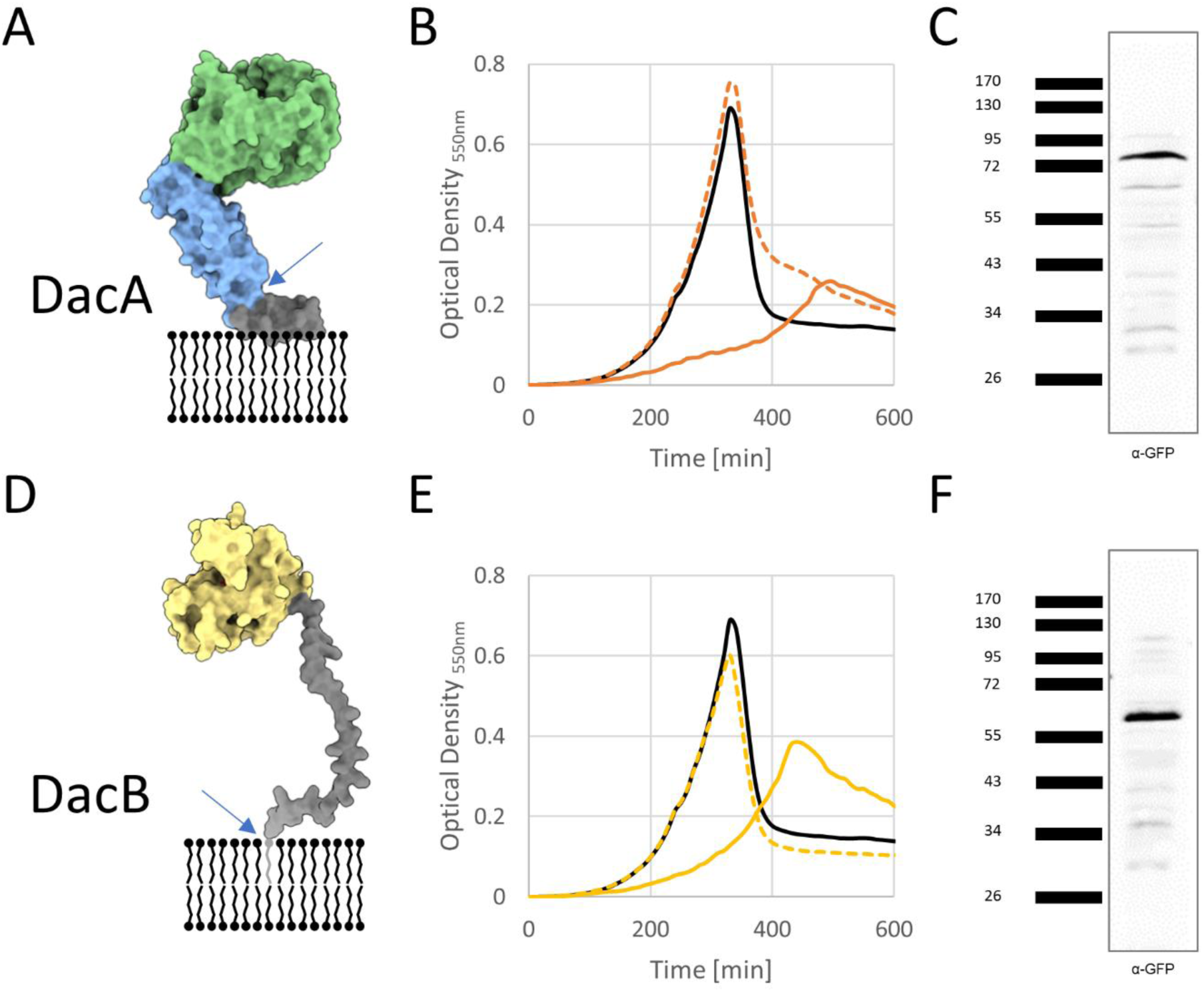
(A) Schematic model of DacA (Q8DQ99). The N-terminal amphipathic helix is shown in dark gray, the globular domain that could act as a pedestal is shown in blue and the domain containing the active site is shown in green. The blue arrow marks the insertion site of sfGFP (B) Representative growth curves of the WT (solid line, black), the ΔdacA mutant (solid line, orange) and the dacA-sfGfp (dashed line, orange) strain. (C) Western blot analysis of the DacA-sfGfp fusion protein constructs detected with an anti-GFP antibody. (D) Schematic model of DacB (Q8DQQ1). The N-terminal disordered region is shown in dark gray and the C-terminal catalytic domain is shown in yellow. The blue arrow marks the insertion site of sfGFP. (E) Representative growth curves of the WT (solid line, black), the ΔdacB mutant (solid line, yellow) and the sfGfp-dacB (dashed line, yellow) strain. (F) Western blot analysis of the sfGfp-DacB fusion protein constructs detected with an anti-GFP antibody.

**Video 1:** A representative time-lapse series of images showing the localization of mKate2-FtsA and the interface between the new and the old poles (aka cell equators) as the cell elongates at the single cell level by time-lapse microscopy. The cell equators can be visualized using a long pulse of the fluorescent D-amino acid (FDAA) followed by a chased experiment performed under the microscope. Pictures were taken every 3 min. The phase contrast, mKate-channel, the HADA-channel and the merged between the mKate and the HADA channels images are shown. The scale bar represents 1 µm.

**Video 2:** A representative time-lapse series of images showing the localization of FtsZ-mGfp and HlpA-mKate (as a proxy for the nucleoid) as the cell elongates at the single cell level by time-lapse microscopy. Pictures were taken every 2 min. The merged between the mKate or the mGfp channels and the phase contrast images are shown. The scale bar represents 1 µm.

**Video 3:** A representative time-lapse series of images showing the localization of mGfp-MapZ as the cell elongates at the single cell level by time-lapse microscopy in a *ΔdacB* mutant cell. Pictures were taken every 2 min. The mGfp channel and the merge between the mGfp channel and the phase contrast images are shown. The scale bar represents 1 µm.

**Table S1:**
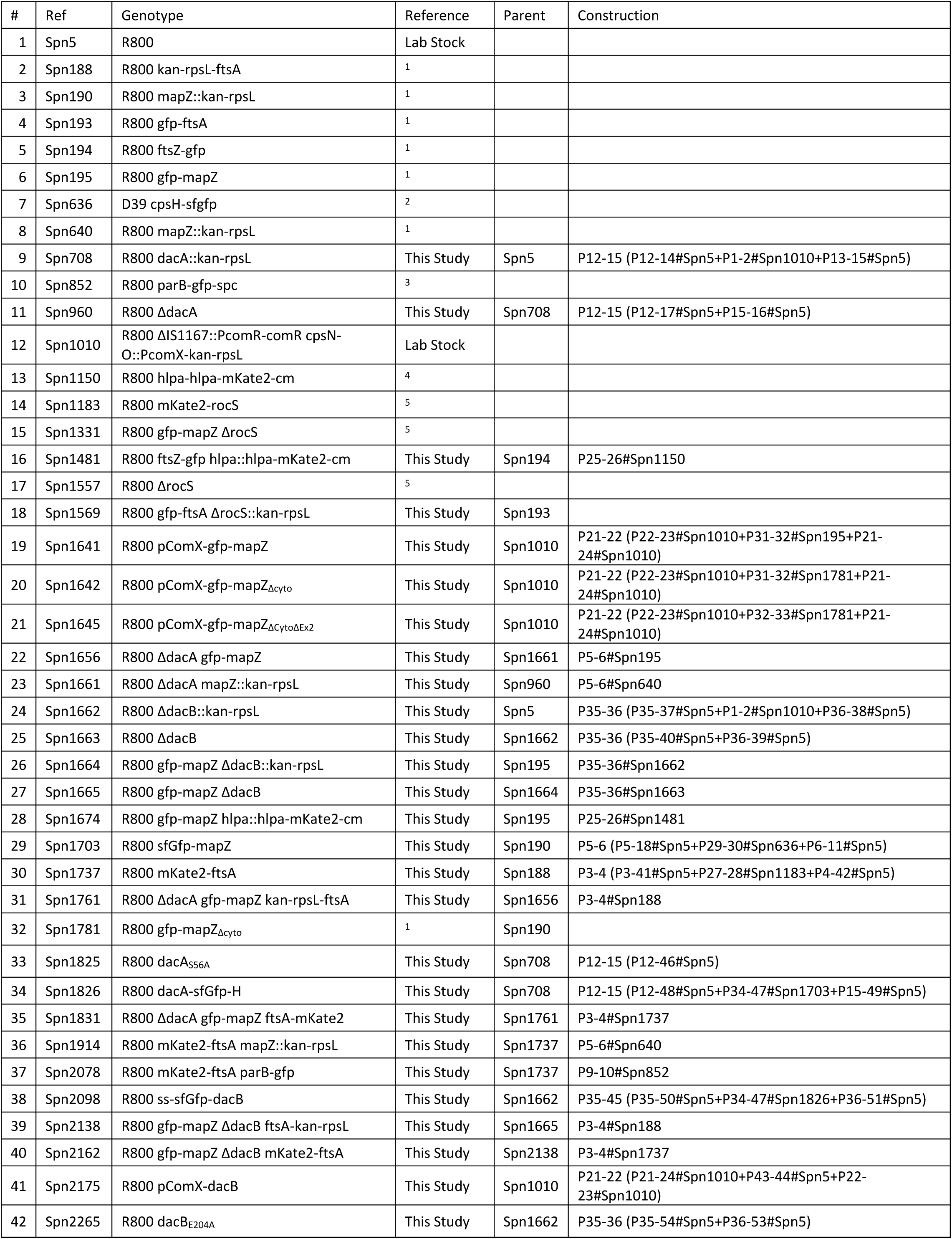
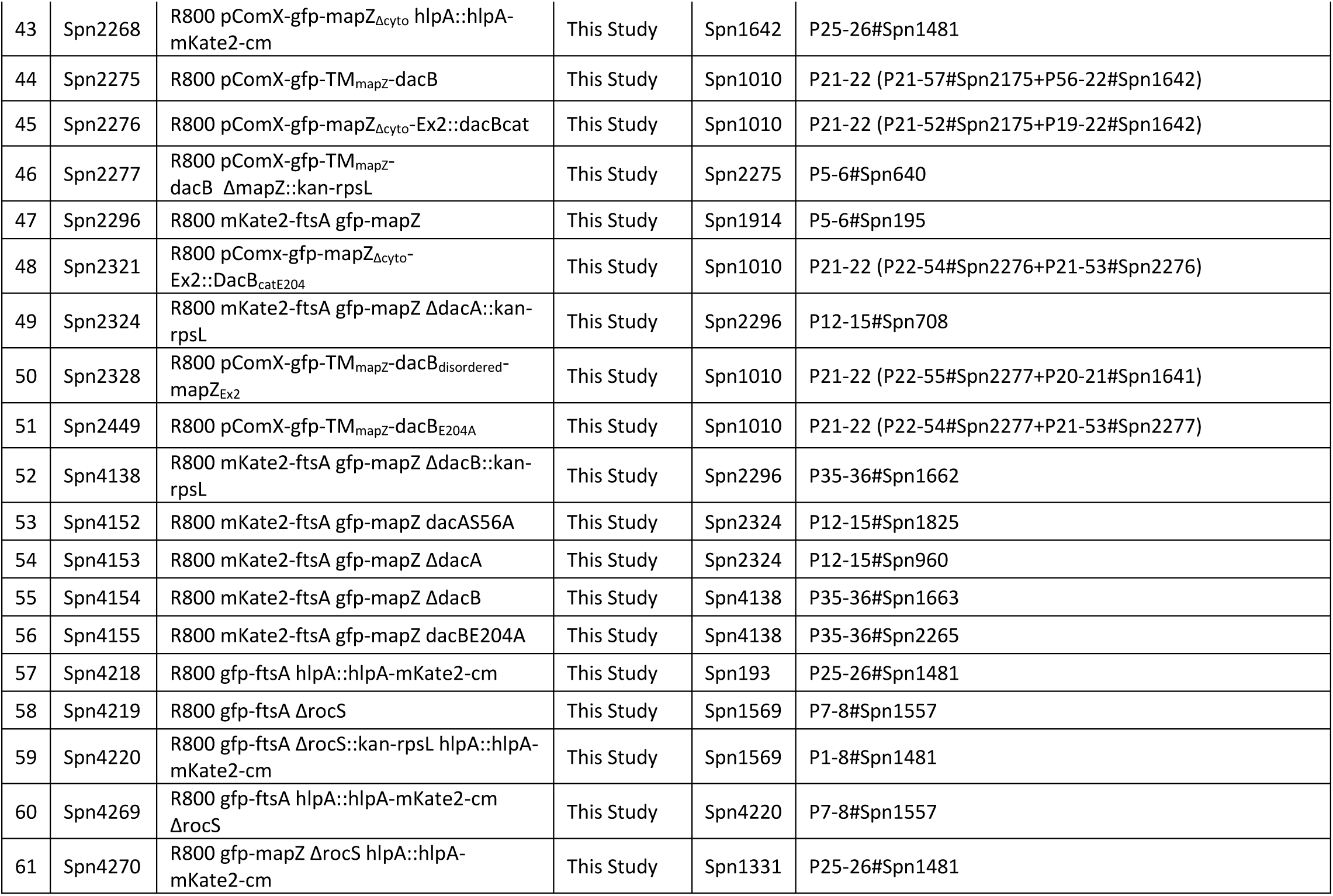
Strains List & Construction.

**Table S2:**
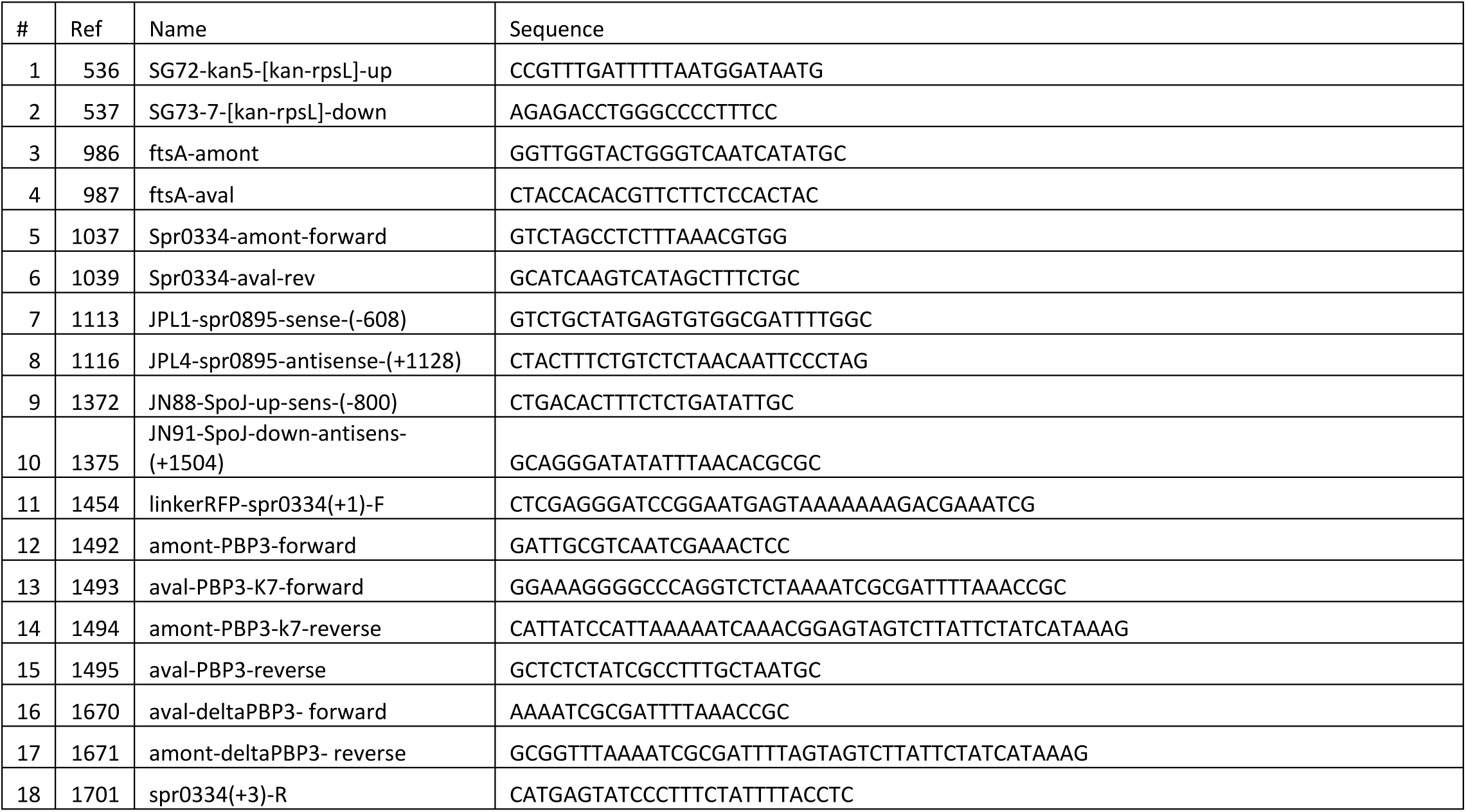
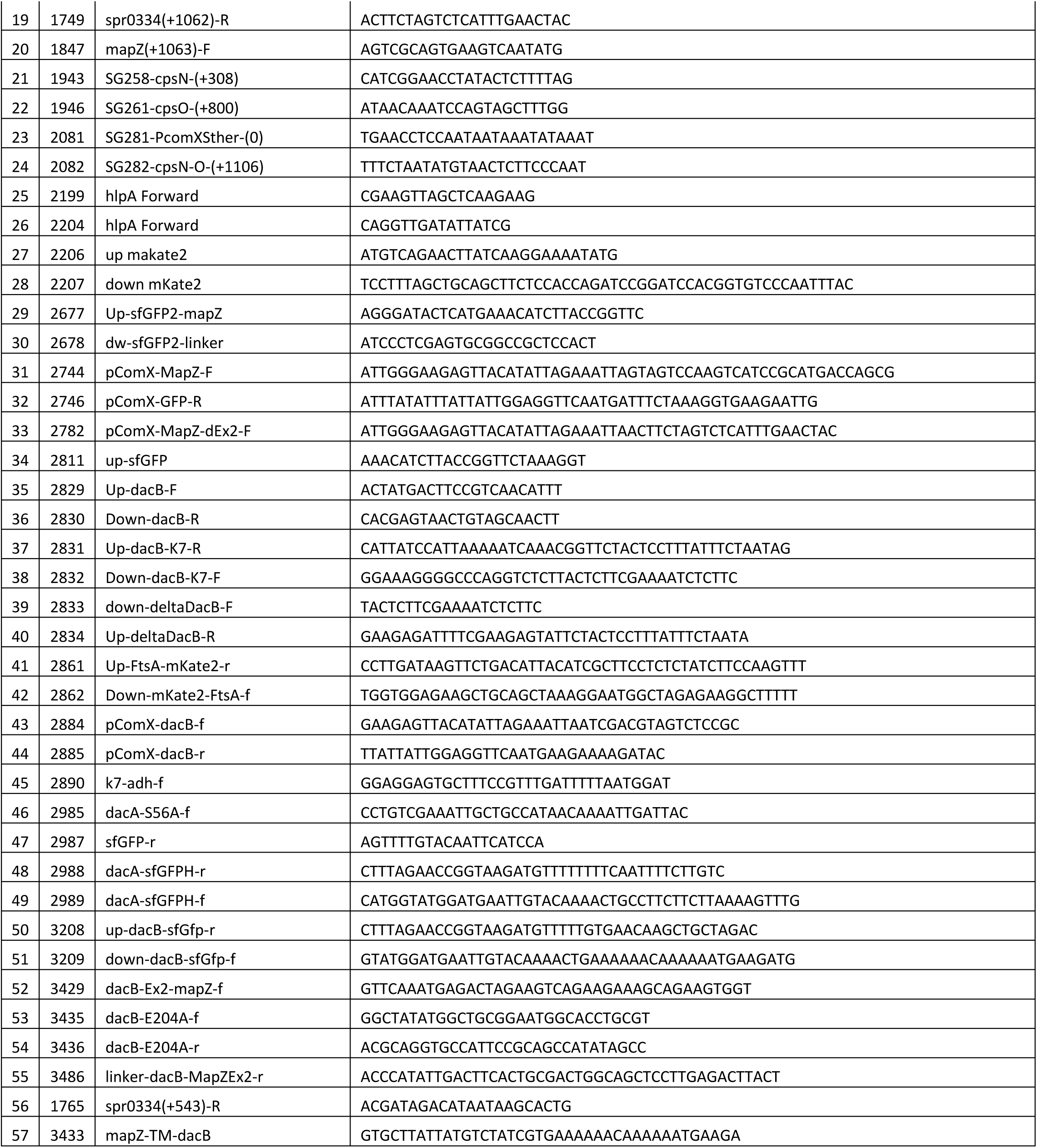
Primers list.

## References

1. Haeusser, D. P. & Margolin, W. Splitsville: structural and functional insights into the dynamic bacterial Z ring. Nat. Rev. Microbiol. 14, 305–319 (2016).

2. Bramkamp, M. & van Baarle, S. Division site selection in rod-shaped bacteria. Curr. Opin. Microbiol. 12, 683–688 (2009).

3. Rowlett, V. W. & Margolin, W. The Min system and other nucleoid-independent regulators of Z ring positioning. Front. Microbiol. 6, 478 (2015).

4. Adams, D. W., Wu, L. J. & Errington, J. Cell cycle regulation by the bacterial nucleoid. Curr. Opin. Microbiol. 22, 94–101 (2014).

5. Thanbichler, M. & Shapiro, L. MipZ, a spatial regulator coordinating chromosome segregation with cell division in Caulobacter. Cell 126, 147–162 (2006).

6. Willemse, J., Borst, J. W., de Waal, E., Bisseling, T. & van Wezel, G. P. Positive control of cell division: FtsZ is recruited by SsgB during sporulation of Streptomyces. Genes Dev. 25, 89–99 (2011).

7. Treuner-Lange, A. et al. PomZ, a ParA-like protein, regulates Z-ring formation and cell division in Myxococcus xanthus. Mol. Microbiol. 87, 235–253 (2013).

8. Ramos-León, F. et al. PcdA promotes orthogonal division plane selection in Staphylococcus aureus. Nat. Microbiol. 9, 2997–3012 (2024).

9. Fleurie, A. et al. MapZ beacons the division sites and positions FtsZ-rings in Streptococcus pneumoniae. Nature 516, 259–262 (2014).

10. Holečková, N. et al. LocZ is a new cell division protein involved in proper septum placement in Streptococcus pneumoniae. mBio 6, e01700–01714 (2014).

11. van Raaphorst, R., Kjos, M. & Veening, J.-W. Chromosome segregation drives division site selection in Streptococcus pneumoniae. Proc. Natl. Acad. Sci. 114, E5959–E5968 (2017).

12. Manuse, S. et al. Structure–function analysis of the extracellular domain of the pneumococcal cell division site positioning protein MapZ. Nat. Commun. 7, 12071 (2016).

13. Perez, A. J. et al. Movement dynamics of divisome proteins and PBP2x:FtsW in cells of Streptococcus pneumoniae. Proc. Natl. Acad. Sci. U. S. A. 116, 3211–3220 (2019).

14. Hosek, T. et al. Structural features of the interaction of MapZ with FtsZ and membranes in Streptococcus pneumoniae. Sci. Rep. 10, 4051 (2020).

15. Li, Y. et al. MapZ Forms a Stable Ring Structure That Acts As a Nanotrack for FtsZ Treadmilling in Streptococcus mutans. ACS Nano 12, 6137–6146 (2018).

16. Massidda, O., Nováková, L. & Vollmer, W. From models to pathogens: how much have we learned about Streptococcus pneumoniae cell division? Environ. Microbiol. 15, 3133–3157 (2013).

17. Vollmer, W., Massidda, O. & Tomasz, A. The Cell Wall of Streptococcus pneumoniae. Microbiol. Spectr. 7, 10.1128/microbiolspec.gpp3-0018–2018 (2019).

18. Ducret, A., Quardokus, E. M. & Brun, Y. V. MicrobeJ, a tool for high throughput bacterial cell detection and quantitative analysis. Nat. Microbiol. 1, 16077 (2016).

19. Mercy, C. et al. RocS drives chromosome segregation and nucleoid protection in Streptococcus pneumoniae. Nat. Microbiol. 4, 1661–1670 (2019).

20. Raghunathan, S., Chimthanawala, A., Krishna, S., Vecchiarelli, A. G. & Badrinarayanan, A. Asymmetric chromosome segregation and cell division in DNA damage-induced bacterial filaments. Mol. Biol. Cell 31, 2920 (2020).

21. Abdullah, M. R. et al. Structure of the pneumococcal l,d-carboxypeptidase DacB and pathophysiological effects of disabled cell wall hydrolases DacA and DacB. Mol. Microbiol. 93, 1183–1206 (2014).

22. Barendt, S. M., Sham, L.-T. & Winkler, M. E. Characterization of Mutants Deficient in the l,d-Carboxypeptidase (DacB) and WalRK (VicRK) Regulon, Involved in Peptidoglycan Maturation of Streptococcus pneumoniae Serotype 2 Strain D39▿. J. Bacteriol. 193, 2290–2300 (2011).

23. Morlot, C. et al. Crystal Structure of a Peptidoglycan Synthesis Regulatory Factor (PBP3) from Streptococcus pneumoniae*. J. Biol. Chem. 280, 15984–15991 (2005).

24. Morlot, C., Noirclerc-Savoye, M., Zapun, A., Dideberg, O. & Vernet, T. The D,D-carboxypeptidase PBP3 organizes the division process of Streptococcus pneumoniae. Mol. Microbiol. 51, 1641–1648 (2004).

25. Schuster, C., Dobrinski, B. & Hakenbeck, R. Unusual septum formation in Streptococcus pneumoniae mutants with an alteration in the D,D-carboxypeptidase penicillin-binding protein 3. J. Bacteriol. 172, 6499–6505 (1990).

26. Briggs, N. S., Bruce, K. E., Naskar, S., Winkler, M. E. & Roper, D. I. The Pneumococcal Divisome: Dynamic Control of Streptococcus pneumoniae Cell Division. Front. Microbiol. 12, (2021).

27. Zhang, J., Yang, Y.-H., Jiang, Y.-L., Zhou, C.-Z. & Chen, Y. Structural and biochemical analyses of the Streptococcus pneumoniae l,d-carboxypeptidase DacB. Acta Crystallogr. D Biol. Crystallogr. 71, 283–292 (2015).

28. Kysela, D. T., Brown, P. J., Huang, K. C. & Brun, Y. V. Biological Consequences and Advantages of Asymmetric Bacterial Growth. Annu. Rev. Microbiol. 67, 417 (2013).

29. Monahan, L. G., Liew, A. T., Bottomley, A. L. & Harry, E. J. Division site positioning in bacteria: one size does not fit all. Front. Microbiol. 5, (2014).

30. Egan, A. J. F., Errington, J. & Vollmer, W. Regulation of peptidoglycan synthesis and remodelling. Nat. Rev. Microbiol. 18, 446–460 (2020).

31. Cho, H. et al. Bacterial cell wall biogenesis is mediated by SEDS and PBP polymerase families functioning semi-autonomously. Nat. Microbiol. 1, 1–8 (2016).

32. Trouve, J. et al. Nanoscale dynamics of peptidoglycan assembly during the cell cycle of Streptococcus pneumoniae. Curr. Biol. CB 31, 2844–2856.e6 (2021).

33. Perez, A. J. et al. Organization of Peptidoglycan Synthesis in Nodes and Separate Rings at Different Stages of Cell Division of Streptococcus pneumoniae. Mol. Microbiol. 115, 1152 (2020).

34. Perez, A. J. et al. Elongasome core proteins and class A PBP1a display zonal, processive movement at the midcell of Streptococcus pneumoniae. Proc. Natl. Acad. Sci. U. S. A. 121, e2401831121 (2024).

35. Sung, C. K., Li, H., Claverys, J. P. & Morrison, D. A. An rpsL Cassette, Janus, for Gene Replacement through Negative Selection in Streptococcus pneumoniae. Appl. Environ. Microbiol. 67, 5190–5196 (2001).

## References

1. Fleurie, A. et al. MapZ beacons the division sites and positions FtsZ-rings in Streptococcus pneumoniae. Nature 516, 259–262 (2014).

2. Nourikyan, J. et al. Autophosphorylation of the Bacterial Tyrosine-Kinase CpsD Connects Capsule Synthesis with the Cell Cycle in Streptococcus pneumoniae. PLoS Genet. 11, e1005518 (2015).

3. van Raaphorst, R., Kjos, M. & Veening, J.-W. Chromosome segregation drives division site selection in Streptococcus pneumoniae. Proc. Natl. Acad. Sci. 114, E5959–E5968 (2017).

4. Kjos, M. & Veening, J.-W. Tracking of chromosome dynamics in live Streptococcus pneumoniae reveals that transcription promotes chromosome segregation. Mol. Microbiol. 91, 1088–1105 (2014).

5. Mercy, C. et al. RocS drives chromosome segregation and nucleoid protection in Streptococcus pneumoniae. Nat. Microbiol. 4, 1661–1670 (2019).

